# Robustness and management performance of MSY reference points derived from the hockey-stick stock-recruitment model under structural uncertainty

**DOI:** 10.64898/2026.03.27.714336

**Authors:** Momoko Ichinokawa, Hiroshi Okamura

**Affiliations:** Fisheries Resources Institute, Japan Fisheries Research and Education Agency, 2-12-4 Fukuura, Kanazawa-ku, Yokohama, Kanagawa 236-8648 Japan; School of Data Science, Yokohama City University, 22-2 Seto, Kanazawa-ku, Yokohama, Kanagawa 236-0027 Japan

**Keywords:** stock-recruitment relationship, hockey-stick, segmented regression, management strategy evaluation, maximum sustainable yield, precautionary approach, bias variance trade-off

## Abstract

The hockey-stick (HS) stock recruitment relationship (SRR) has been widely used as an empirical alternative to conventional SRRs such as the Beverton–Holt (BH) and Ricker (RI) models. However, the management performance and risks associated with estimating maximum-sustainable-yield (MSY) reference points (RPs) based on HS remain insufficiently understood. This study first defines deterministic and stochastic MSY RPs under the HS model and provides an overview of their properties. We then conduct simulation experiments to investigate the bias and management consequences that arise when MSY RPs are estimated from the HS model (HS-derived MSY RPs) rather than from the true SRR (e.g., BH) across a range of biological and stochastic parameters, with particular focus on scenarios with insufficient data contrast. Our results show that HS-derived MSY RPs tend to exhibit higher bias but lower variance than MSY RPs derived from the true SRR. Management strategy evaluation simulations further reveal that management procedures combining HS-derived MSY RPs with adaptive model learning and some precautionary measures gradually reduce this bias and achieve average spawning biomass and yield that are comparable to those obtained under management based on the true BH SRR. We also show that the management effectiveness of the precautionary measures depends on life-history traits and recruitment variability. These findings indicate that although HS-derived MSY RPs may be biased and require cautious use, combining them with appropriate precautionary measures allows management to remain robust while limiting variability and yield losses. This broadens the range of management options that are available for supporting sustainable fisheries management.

## Introduction

Understanding the productivity of fish stocks and adjusting fishing practices accordingly is fundamental for achieving sustainable fisheries management. Managing fisheries resources to maximize their productivity, which is defined as maximum sustainable yield (MSY), has been established as a central goal of fisheries management under the United Nations Convention on the Law of the Sea and many fisheries management organizations (Hilborn and Stokes 2010). The concept of MSY is derived from models in which the productivity of a fish stock is represented as a concave function of population abundance. In age-structured population dynamics, the relationship between productivity and population abundance is largely determined by the relationship between recruitment and spawning stock biomass (stock-recruitment relationship, SRR). Accurate estimation and prediction of the SRR is therefore essential for quantifying stock productivity and for determining key management reference points such as MSY.

However, inferring the SRR accurately is widely recognized as one of the most challenging issues in stock assessment and management (Walters and Martell 2004). Simulation studies have shown that reliable estimation of SRR parameters requires both sufficient contrast in spawning stock biomass and correct model specification (Lee et al. 2014).

The hockey-stick (HS), segmented regression SRR (Clark et al. 1985, Mesnil et al. 2010) has been proposed as an alternative SRR when traditional models such as the Ricker (RI) (Ricker 1954) and Beverton–Holt (BH) (Beverton and Holt 1957) models are difficult to apply because of insufficient data contrast (ICES 2022, FRA 2025a). An important advantage of HS in a management context is that it avoids the extrapolation of recruitment beyond the range of historically observed values, as such extrapolation can lead to unrealistic MSY reference point (RP) estimates (Clark et al. 1985, Ichinokawa et al. 2017). HS can also avoid predicting unrealistically high numbers of recruits per spawner at very low population sizes, as predicted by BH and RI (Barrowman and Myers 2000). Although HS was not originally developed from biological theory (Walters and Martell 2004), recent work has provided an ecological interpretation for SRR shapes, similar to that represented by HS (Maunder 2022).

In practice, situations with insufficient data contrast to reliably determine the SRR are common in stock assessments. For example, Silvar-Viladomiu et al. (2022) reported that more than half of data-rich stocks with full analytical assessments (category-1) in the International Council for the Exploration of the Sea (ICES) are classified as stocks for which no clear evidence of impaired recruitment has been observed. Similarly, Ichinokawa et al. (2017) showed that no single SRR could be statistically selected among three candidate SR models for 20 of the 26 Japanese stocks. Under such circumstances, the HS model is often used in stock assessments to derive management reference points. According to recent stock assessments, HS (referred to as segmented regression by ICES) plays a role in determining management RPs in approximately half of the 74 category-1 stocks assessed by ICES (https://standardgraphs.ices.dk/stockList.aspx, accessed December 2025). In Japan, the HS model was explicitly used for determining MSY RPs in 21 of 31 Japanese domestic stocks in the 2024 stock assessment (https://abchan.fra.go.jp/hyouka/doc2024/, accessed January 2026). These examples illustrate the widespread use of HS as a practical alternative SRR in fisheries management.

Under the HS model, management RPs are largely determined by the position of the change point, i.e., the historical range of stock abundance, rather than the intrinsic productivity of the stock. For example, when recruitment appears to be independent of spawning biomass, the minimum historical spawning biomass becomes the change point. Conversely, when recruitment increases approximately linearly with spawning biomass, the maximum historical spawning biomass becomes the change point. Such cases often arise when data contrast is insufficient, for example due to a short assessment period or relatively stable stock abundance under consistent fishing pressure (Ichinokawa et al., 2017). As a result, HS-derived MSY reference points may be biased and potentially compromise sustainable fisheries management.

The potential advantages and risks of using MSY RPs derived from the HS model, as well as the theoretical basis for their definition, have not yet been fully evaluated or investigated. Mesnil et al. (2010) suggested that the deterministic F_MSY_ (the fishing mortality rate when the average total yield is maximized under deterministic and equilibrium population dynamics) obtained when applying HS SRR corresponds to either F_max_ (the fishing mortality rate when the total yield per recruitment is maximized) or F_crash_ (the fishing mortality rate above which the deterministic population dynamics crash). Recent studies calculated MSY RPs under stochastic conditions and demonstrated that these values differ from those derived under deterministic assumptions when HS is used (Ichinokawa et al. 2017, Okamura et al. 2020, Okamura et al. 2025). Okamura et al. (2020) further reported that stochastic HS-derived MSY RPs can provide robust management performance. Despite these advances, a comprehensive understanding of how life history parameters and structural misspecification influence HS-derived MSY reference points and their management consequences remains lacking.

This paper addresses the following questions.

1. How can MSY RPs be defined and calculated under the HS model for both deterministic and stochastic population dynamics?
2. When HS is applied instead of the true SRR, such as BH or RI, how and to what extent do HS-derived MSY RPs deviate from the true MSY RPs?
3. How can HS-derived MSY RPs be incorporated into management procedures for achieving sustainable fisheries management?

We focus particularly on situations with insufficient data contrast, where reliable estimation of SRR parameters is difficult. Firstly, utilizing simulations of age-structured population dynamics with BH, RI, and HS models, we evaluated MSY RPs under a wide range of life history parameters and recruitment variability conditions. We then examined the management performance achieved when HS-derived MSY RPs were used in harvest control rules under various stock depletion scenarios. By addressing these questions, we clarify the properties of HS-derived MSY RPs and provide pragmatic guidance for their appropriate use in fisheries management.

## Materials and methods

### Structure of the simulation tests for addressing the three main objectives

This study was hierarchically structured to address the three main objectives (Fig. 1). In the first analysis, (“A. Simulation design and true MSY RPs”), simulations were conducted using the stochastic population dynamics model to estimate MSY RPs under a variety of biological parameters and SRRs: all combinations of three SRRs (HS, BH, and RI), four sets of plausible life history parameters, a range of steepness values, and three levels of recruitment variability. In the second analysis (“B. Estimation experiments”), we examined the relative bias and variances of MSY RP estimates obtained when the HS SRR was applied to the simulated data generated from BH under scenarios with varying stock depletion levels without data contrast. In the third analysis (“C. Management strategy evaluation”), we extended the simulation period starting from step B to evaluate management performance under procedures based on the estimated HS-derived MSY RPs, as a simplified management strategy evaluation (MSE).

**Fig. 1.**
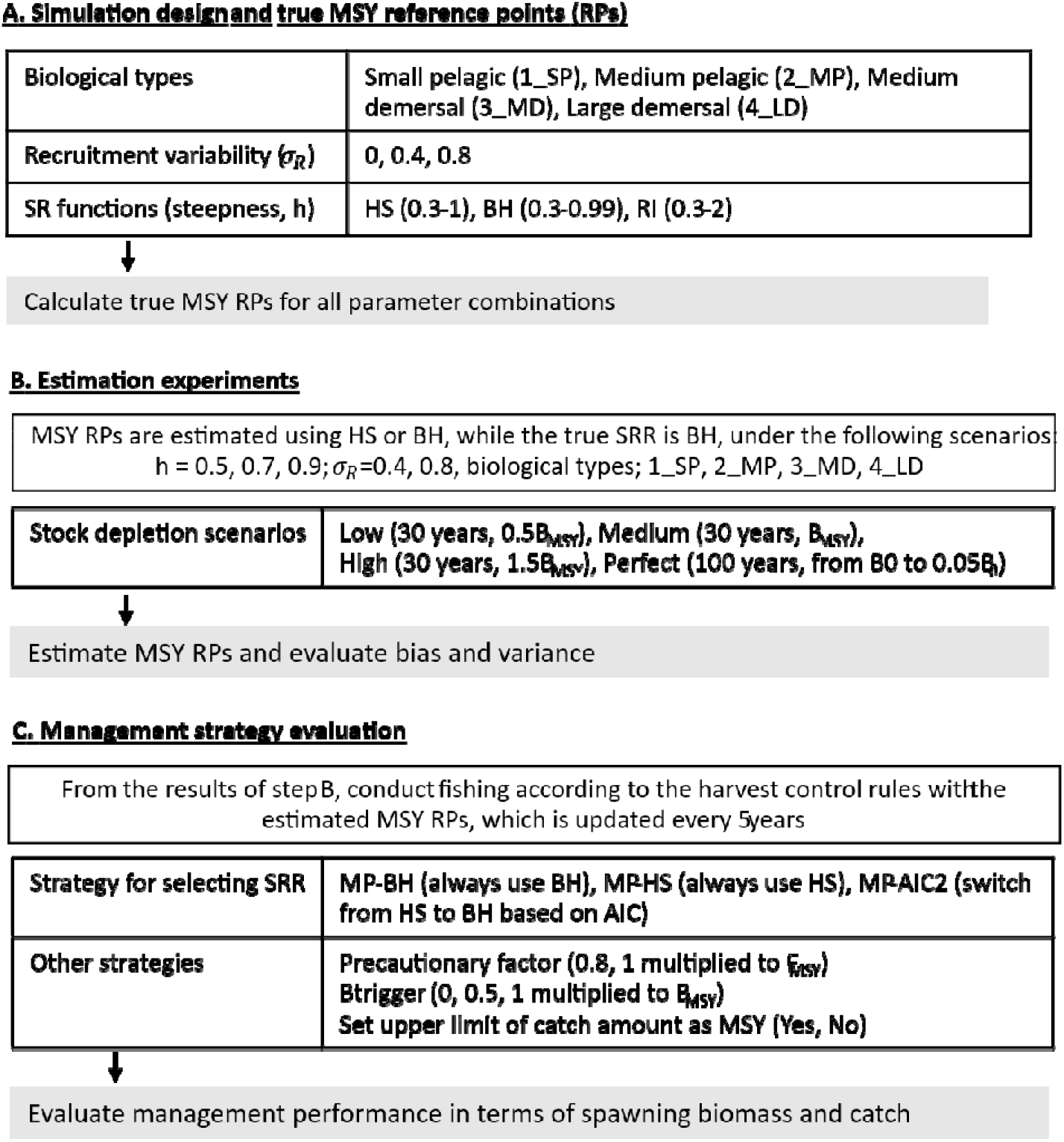
Overview of the simulation design, estimation experiment, and management strategy evaluation.

### Population dynamics

The age-structured population dynamics used in this analysis are indexed by year *y*, age *a*, and simulation iteration *k*, and variables are denoted with these subscripts as needed. We categorized the simulation period as the burn-in period before stock assessment (*T*_0_ < *y* < *T*_1_ stock assessment period (*T*_1_ ≤ *y* < *T*_2_), and stock management period (*T*_2_ ≤ *y* ≤ *T*_3_). The number of fish 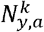 in the population is given by:

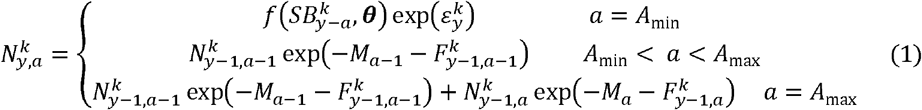

where *M*_*a*_ is the natural mortality rate; 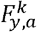 is the fishing mortality coefficient; *f* is the SR function of BH, RI or HS (Eqs. A3–A5 in Appendix A, respectively); 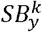 is the spawning biomass; ***θ*** is a parameter vector of the SR function; 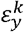 is the process error that is randomly generated from N (− 0.5*σ*^2^, *σ*^2^); *A*_min_ is the recruitment age; and *A*_max_ is the plus group age. The spawning biomass 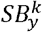 is defined as 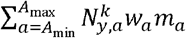 where *w*_*a*_ and *m*_*a*_ are the mean weight and maturity rate at age, respectively. The fishing mortality coefficient 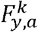 was calculated as the multiplier of *S*_*a*_ (the selectivity at age *a*, with a maximum of 1) and 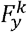 (the fishing mortality coefficient at the fully selected age and year *y*). 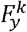 was determined according to the stock depletion scenario before and during the stock assessment period (generally *y* = 1,2, … *T*_2_ − 1) and harvest control rule in the management period (*y* =*T*_2_, …,*T*_3_). Total catch weight in year *y* and age 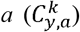 was calculated from Baranov catch equation (Eq. 6 in Appendix A). More detailed equations and simulation settings are available in Appendix A.

The biological parameters *M*_*a*_, *m*_*a*_, and *w*_*a*_ were derived from four representative Japanese species differing in body size, longevity, and habitat (pelagic or demersal): a small pelagic species (Japanese sardine, 1_SP; Furuichi et al. 2024), a medium-sized pelagic species (chub mackerel, 2_MP; Yukami et al. 2024), a medium-sized demersal species (walleye pollock, 3_MD; Chimura et al. 2024), and a large demersal species (Japanese flounder, 4_LD; Masubuchi et al. 2024). The biological parameters are presented in Fig. B of Appendix B. In the base case, the fishing selectivity rate at age *S*_*a*_ was assumed to be equal to the maturity rate at age, but values less than 0.1 were replaced by 0.1 to avoid F_MAX_ become infinity in some species. For the sensitivity analysis, we used the selectivity patterns averaged over the stock assessment period for each stock (Fig. B-d).

The stochastic simulations were implemented in R (R Development Core Team, 2024), and the code is available at https://github.com/ichimomo/HS-simulation.

#### A. True MSY RPs

In step A, we calculated MSY RPs with a variety of life histories and SRR parameters under three SRRs of HS, BH and RI. In this study, we calculated the MSY RPs based on the stochastic simulations described by Eq. (1). We defined the MSY as the maximized equilibrium average catch weight across 1,000 stochastic simulations under constant fishing selectivity (Appendix A). SB_MSY_ and F_MSY_ were defined as the average spawning biomass and fishing mortality rate, respectively, when the MSY was obtained. Similarly, SB_0_ was defined as the equilibrium average of the spawning biomass in the absence of fishing.

The SRR parameters were determined from the strength of the recruitment compensation expressed by the steepness parameter (*h*). The parameter *h* is defined as the fraction of recruitment from an unfished (virgin) population when the spawning biomass is reduced to 20% of SB_0_ (Mangel et al. 2009). For the BH SRR, *h* was varied from 0.3 to 0.95 in increments of 0.05, and 0.99 was also included. For the RI SRR, *h* was varied from 0.3 to 2.0 in increments of 0.1. For the HS SRR, *h* was defined based on the spawning biomass of the break point of HS (SB_HS_) as 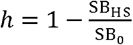 (Punt et al. 2014) and varied from 0.3 to 0.95 in increments of 0.05. All the variations exhibited by the SR functions that were used to calculate MSY RPs are illustrated in Fig. A1. The recruitment variability (*σ*) was set to 0, 0.4, and 0.8.

#### B. Evaluation of the biases and variances of MSY RPs

In step B, we estimated the parameters of the SR functions and the corresponding MSY RPs using the SR data generated from the simulated population dynamics under specified depletion scenarios. In this simulation, we assumed that the true values of recruitment and spawning biomass were known without time lags and were not subject to stock assessment or observation errors. This assumption was made to evaluate the pure effects of the biases of the SR parameters and the misspecification of the SR model under SR data that were limited by the sampling period (31 years, *T*_1_ = 20, *T*_2_ = 50) and the lack of data contrast (i.e., no substantial stock abundance variations), even when the SR data were sufficiently accurate.

The historical stock trends (stock depletion scenarios) were controlled using the parameter *P*_*y*_, which was defined as the ratio of the spawning biomass to SB_MSY_ in year *y*. Fishing mortality was determined to satisfy the conditions specified by *P*_*y*_ in each scenario under deterministic population dynamics, and the same fishing mortality was then applied in stochastic simulations incorporating random recruitment variability. We considered three realistic scenarios without data contrast. In these scenarios, the stock level was held constant (i.e., 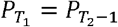) by applying a constant fishing mortality rate. The SR data were collected from *y* = *T*_1_ to *T*_2_ − 2. The simulated constant stock levels were 0.5SB_MSY_ (low: 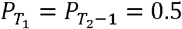), 1.0SB_MSY_ (middle: 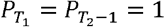), and 1.5SB_MSY_ (high: 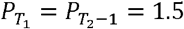) (Fig. A2). For comparison purposes, we also considered an unrealistic “perfect” scenario with sufficient data contrast. In the perfect scenario, the SR data were collected from *y* = 20 (with the spawning biomass at SB_0_) to *y* = 100 (with *P*_100_ = 0.05), assuming a linear increase in fishing mortality. The biological parameters assumed in those simulations were essentially the same as those used in step A. The results for 1_SP with recruitment variability (*σ* > 0) are presented as the main analysis, while those for the other species are provided in the Supplementary materials.

As the estimation experiments in step B, the SR data sampled from the population dynamics simulations were applied to the SR functions to estimate their parameters and MSY RPs as 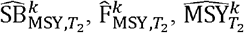, and 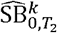 The superscript *k* indicates the *k*-th simulation replicate, and the subscript *T*_2_ indicates the year when the MSY RPs were estimated using the data from *y* = 20 to *T*_2_ − 1. To evaluate bias and variance of the MSY RPs, we calculated relative errors of the estimated MSY RPs. For example, the relative error of SB_MSY_ (RE(SB_MSY,y_)) was calculated as 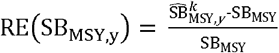, where the true values SB_MSY_ were derived from those calculated in step A.

We focused especially on the situation where BH was the true SRR but HS was applied for the estimation of the MSY RPs (i.e., the expedient application of HS), and we compared these results with those obtained when BH was correctly applied to the simulation data. In the analyses of steps B and C, the RI function was not included in the main simulation scenarios to reduce the number of scenario combinations, but we conducted a sensitivity analysis by replacing BH with RI. The biological parameters used in steps B and C were all combinations of h = 0.5, 0.7, or 0.9; *σ* = 0.4 or 0.8; and four species (1_SP, 2_MP, 3_MD, 4_LD) and number of replicates was 300 (Fig. 1).

#### C. Management strategy evaluation (MSE)

The long-term stochastic simulations were conducted from *y* = 51 to 90 under all combinations among biological parameters, stock depletion scenarios, and harvest control rules (Fig. 1). The harvest control rule (HCR) used to determine 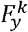 was defined as

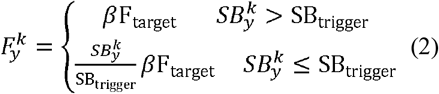

where the precautionary factor *β* was set to 0.8 or 1 and the spawning biomass threshold SB_trigger_ as 0, 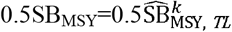 or 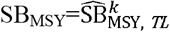, while target fishing mortality (F_target_) was fixed at 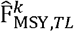, where *TL* denotes the most recent year in which the MSY RPs were updated. The MSY RPs were updated every 5 years to reduce the calculation time as *TL* = 51,56,61, …,86. In addition, the candidate MPs further diverged with respect to the following two aspects: (1) the method for selecting the SR functions that were used to estimate the MSY RPs (SR selection strategy) and (2) the application of a catch cap (capping strategy). With respect to the SR selection strategy, the MSY RPs used as SB_trigger_ and F_target_ were estimated from the following three scenarios: MP-BH (always using BH), MP-HS (always using HS), and MP-AIC (primarily using HS but switching to BH if Akaike Information Criteria (AIC) of BH was smaller than AIC of HS by more than 2 units). For the capping strategy, we considered whether the annual total catch was capped by the estimated MSY 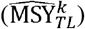 as a precautionary management measure. This was denoted by /C (with capping) or /N (without capping).

While the total number of evaluated MPs was 36 (combinations of two *β*, three Btrigger, three SR selection, and two capping scenarios), we particularly compared the performance among the five MPs: MP-BH/N, MP-HS/C, MP-AIC/N, MP-AIC/N, MP-HS/C under *β* = 0.8 and SBtrigger = SB_MSY_. MP-BH/N (MP-BH without capping) was considered the reference scenario in which the SR function was correctly specified, whereas MP-HS/N (MP-HS without capping) represented the naïve alternative strategy with HS. The three alternative MPs of MP-HS/C (MP-HS with capping), MP-AIC/N (MP-AIC without capping), and MP-AIC/C (MP-AIC with capping) were regarded as advanced strategies in which the concept of adaptive learning and/or precautionary measures was incorporated into the strategy with HS.

We summarised the performance of the five MPs using the following eight statistics (Table). The two statistics for stock abundance were measured as the average ratios of the spawning biomass to the true SB_MSY_ at the 10th and 40th management years after the introduction of management (SB10 and SB40, respectively). We also quantified the stock abundance risk by calculating the probability that the spawning stock biomass remained above 0.5 times the true SB_MSY_ throughout the 10th to 40th management years (SBsafe). For yield performance, we used the average ratios of the catch to the true MSY in the 10th and 40th management years (AC10 and AC40, respectively). Additionally, we assessed the extent of the initial catch reduction at the management introduction year by calculating the average ratio of the total catch in the first management year relative to the catch in the previous year (Cinit). For variability, the average annual variability (AAV) was calculated as the average of |*x*_*y*_ − *x*_*y* − *s*_ | /*x*_*y* − *s*_, where *x*_*y*_ is the value of the metric in year *y* and *s* is the time lag for comparison. The AAV of the total catch during the 10th–40th management years (CAAV) was calculated according to 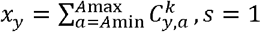, and *y =* 52,52, …,90. The AAV of the estimated SB_MSY_ (SBmsyAAV) was calculated according to 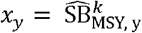, *s* = 5, and *y* = 56,61, …,86. AAV and SBmsyAAV were averaged across all simulation iterations. Since large catch fluctuations and frequent management target level changes are generally undesirable, smaller CAAV and SBmsyAAV values indicate better performance.

**Table.**
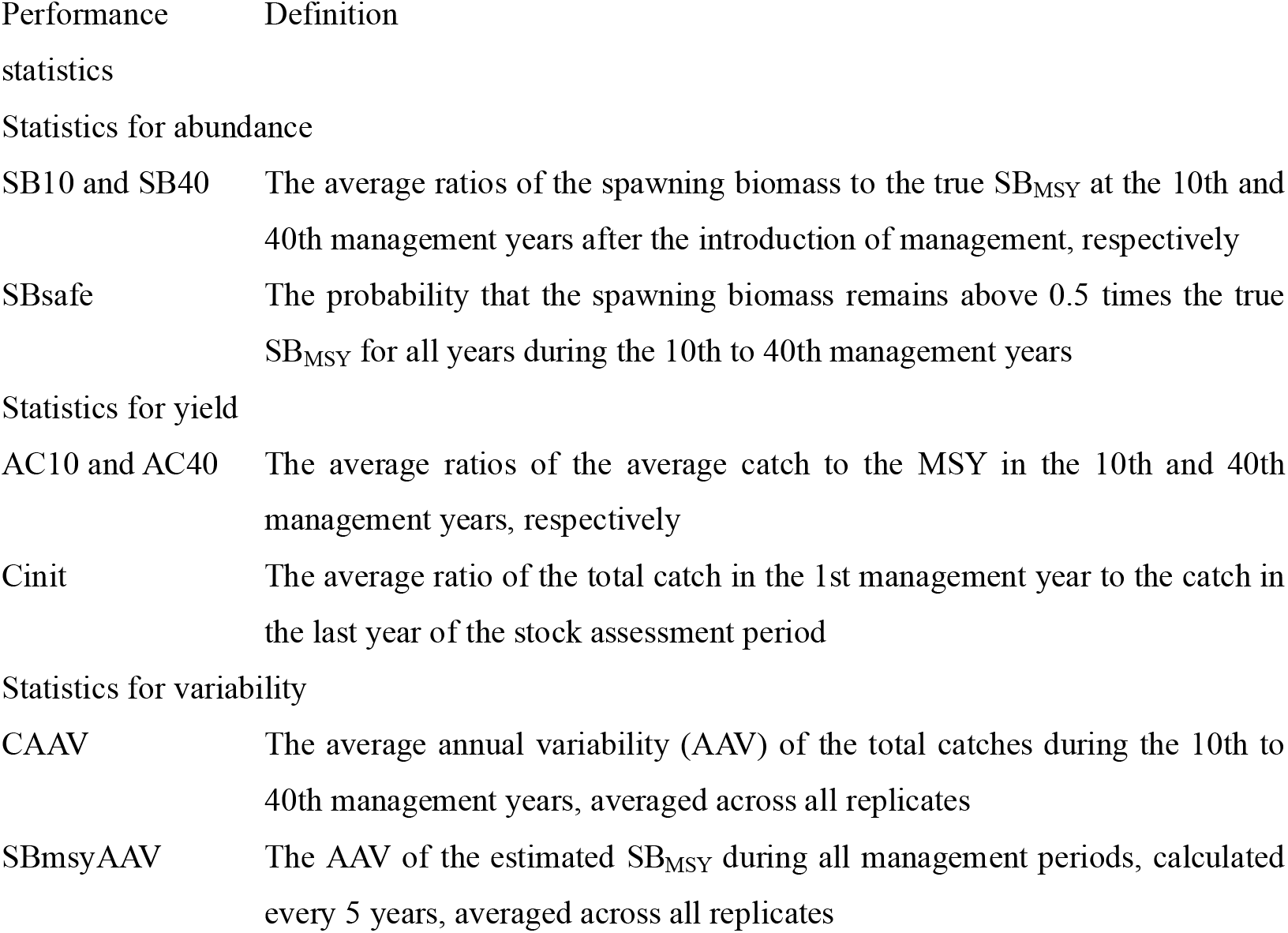
Selected performance statistics.

| Performance | Definition |
| --- | --- |
| SB10 and SB40 | The average ratios of the spawning biomass to the true $SB_{MSY}$ at the 10th and 40th management years after the introduction of management, respectively |
| SBsafe | The probability that the spawning biomass remains above 0.5 times the true $SB_{MSY}$ for all years during the 10th to 40th management years |
| Statistics for yield |  |
| AC10 and AC40 | The average ratios of the average catch to the MSY in the 10th and 40th management years, respectively |
| Cinit | The average ratio of the total catch in the 1st management year to the catch in the last year of the stock assessment period |
| Statistics for variability |  |
| CAAV | The average annual variability (AAV) of the total catches during the 10th to 40th management years, averaged across all replicates |
| SBmsyAAV | The AAV of the estimated $SB_{MSY}$ during all management periods, calculated every 5 years, averaged across all replicates |

To examine which the default precautionary measures most strongly influence management success or failure across species and stock depletion levels, we applied a generalized linear model (GLM) to the full set of simulation results across all 36 MPs. The analysis was conducted for each stock depletion level. For each scenario *i*, defined by a combination of parameter settings, we calculated the number of successful outcomes. We defined two types of management success corresponding to biological and fishery performance: (i) the spawning biomass in the 40th year exceeded the true SB_MSY_ (SB_success_) and (ii) the average catch in years 10 to 40 exceeded 0.9 MSY (Catch_success_). Then, the number of successful outcomes *v*_*i*_ was modelled as follows:

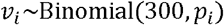

where *p*_*i*_ represents the probability of management success for scenario *i*. A logit link function was used to relate *p*_*i*_ to explanatory variables representing precautionary measures and other biological factors as follows:

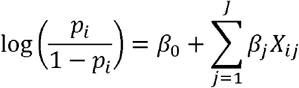

where *X*_ij_ denotes the *j* th explanatory variable for scenario *i* and *β*_*j*_ is the estimated coefficient. All explanatory variables were treated as categorical for *σ* _*R*_ (<u>0.4</u> or 0.8), h (0.5, <u>0.7</u>, or 0.9), SR selection strategies (<u>MP-BH</u>, MP-HS, or MP-AIC), capping (/C or /<u>N</u>), *α* (0.8 or <u>1</u>), and the definition of Btrigger (<u>0</u>, 0.5SBmsy, or SBmsy), where the underlined categories were used as reference levels. The estimated coefficients represent log-odds ratios relative to the reference levels, with positive (negative) values indicating higher (lower) odds than those of the reference while holding the other variables constant. Because the data were generated from simulations and the inclusion of interaction terms reduced the dispersion factor toward 1, interaction effects are likely to be present. To maintain interpretability, however, we present a simplified model including only main effects.

For the sensitivity analyses, we conducted a series of analysis (steps B and C) assuming RI as the true SR relationship. The biological parameters were largely identical to those used in the base case, except that h was set to 0.7, 1.5, or 2, allowing for stock-recruitment dynamics that cannot be represented by the BH model. In addition, MP-BH (always using BH) was replaced with MP-RI (always using RI), and MP-AIC was defined as a strategy in which HS was primarily used but switched to RI if the AIC of RI was smaller than the AIC of HS by more than 2 units. Detailed results are not shown due to space limitations; the results of the GLM analysis are provided in the Appendix.

## Results

### MSY RPs under the HS SR function

While the MSY RPs varied with the SR function type, life history parameters, and steepness (*h*), recruitment variability (*σ*) also substantially influenced the MSY RPs, particularly under the HS function (Fig. 2). A higher *σ* generally resulted in a higher %SPR values corresponding to F_MSY_ (%SPR_MSY_) (i.e., lower F_MSY_ values), higher SB_MSY_ values, and lower MSY values across almost all ranges of *h* under HS, except for SB_MSY_ when *h* was less than 0.5. Although some dependence of the MSY RPs on *σ* was also observed under BH and RI in certain cases (e.g., RI with steepness levels greater than 1 and *σ* = 0.8), the differences were relatively minor. Another characteristic observed under HS was that the curves of MSY RPs as a function of *h* became flatter when *h* was high and *σ* was low. The details of these phenomena are discussed further in the Discussion section.

**Fig. 2.**
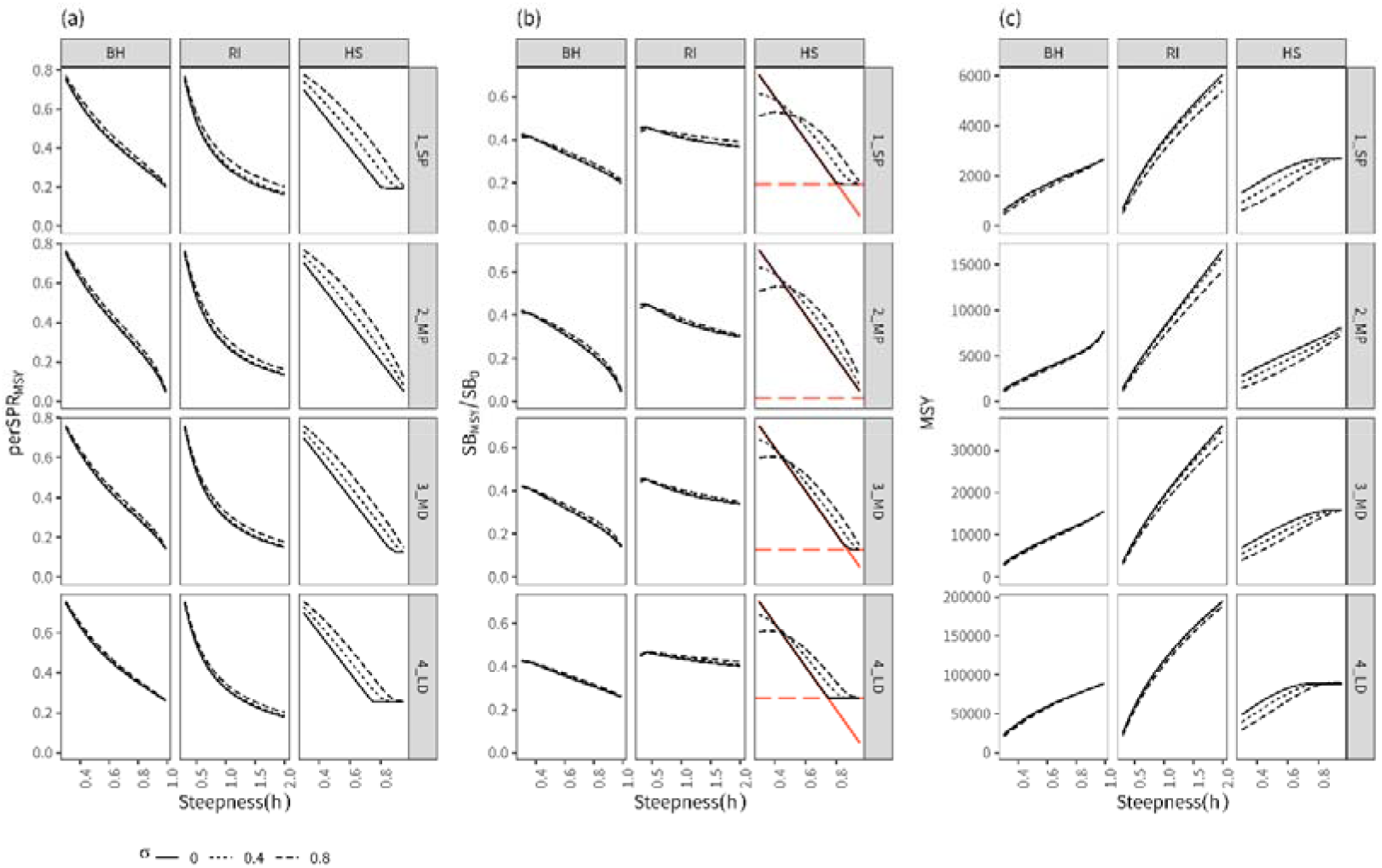
Deterministic (*σ* = 0) and stochastic (*σ* = 0.4 or 0.8) MSY reference points under various types of biological parameters, steepness and SR functions: (a) the %SPR corresponding to the fishing mortality rate when achieving MSY (%SPR_MSY_, perSPR_MSY_), (b) the spawning biomass achieving MSY (SB_MSY_) relative to the unfished biomass (SB_0_), and (c) MSY. The red diagonal broken lines in (b) represent the breakpoint position of the spawning biomass in the HS function (SB_HS_), and the red horizontal lines represent the spawning biomass corresponding to maximum yield per recruit (SB_MAX_); the details are described in the Discussion section. The rows correspond to the species defined in Fig. 1.

We also found that the range of relative SB_MSY_/SB_0_ values when using HS was wider than those observed under BH or RI (Fig. 2). Under HS, the 90th percentile ranges of the distributions of SB_MSY_/SB_0_ across all combinations of biological parameters were from 0.15 to 0.64, with a median of 0.44. In contrast, the ranges were narrower for BH (0.15 to 0.42, with a median of 0.32) and RI (0.32 to 0.46, with a median of 0.41). In contrast, the 5th and 95th percentiles of %SPR_MSY_ were similar across the three SRRs: 0.15 (HS), 0.17 (BH), and 0.16 (RI) for the 5th percentiles and 0.73, 0.75, and 0.75 for the 95th percentiles, respectively. However, the median %SPR was lower for RI (0.28) than for BH (0.42) and HS (0.45), reflecting the inclusion of values of *h* > 1 in the RI function. These results were robust to the assumption of selectivity, as the differences in the medians and 90th percentiles were less than 4%, even when alternative selectivity assumptions based on the historical average for each stock were used.

### Estimation biases induced when misspecifying the SR function

Even when the true SRR was BH and correctly applied, the estimates of F_MSY_, SB_MSY_, and MSY varied and were biased under the “low”, “middle”, and “high” no-data-contrast scenarios, whereas the MSY RPs were estimated with little bias and with relatively small variances under the “perfect” scenario (Fig. 3, Figs. C1-3). The biases observed in the relative errors of F_MSY_ and SB_MSY_ when BH was used under the no-data-contrast scenarios were generally positive and negative, respectively, regardless of the depletion level. The magnitudes of these biases and variances increased as the steepness decreased. The biases and variances exhibited by the relative errors of the MSY tended to be smaller than those observed for F_MSY_ and SB_MSY_.

**Fig. 3.**
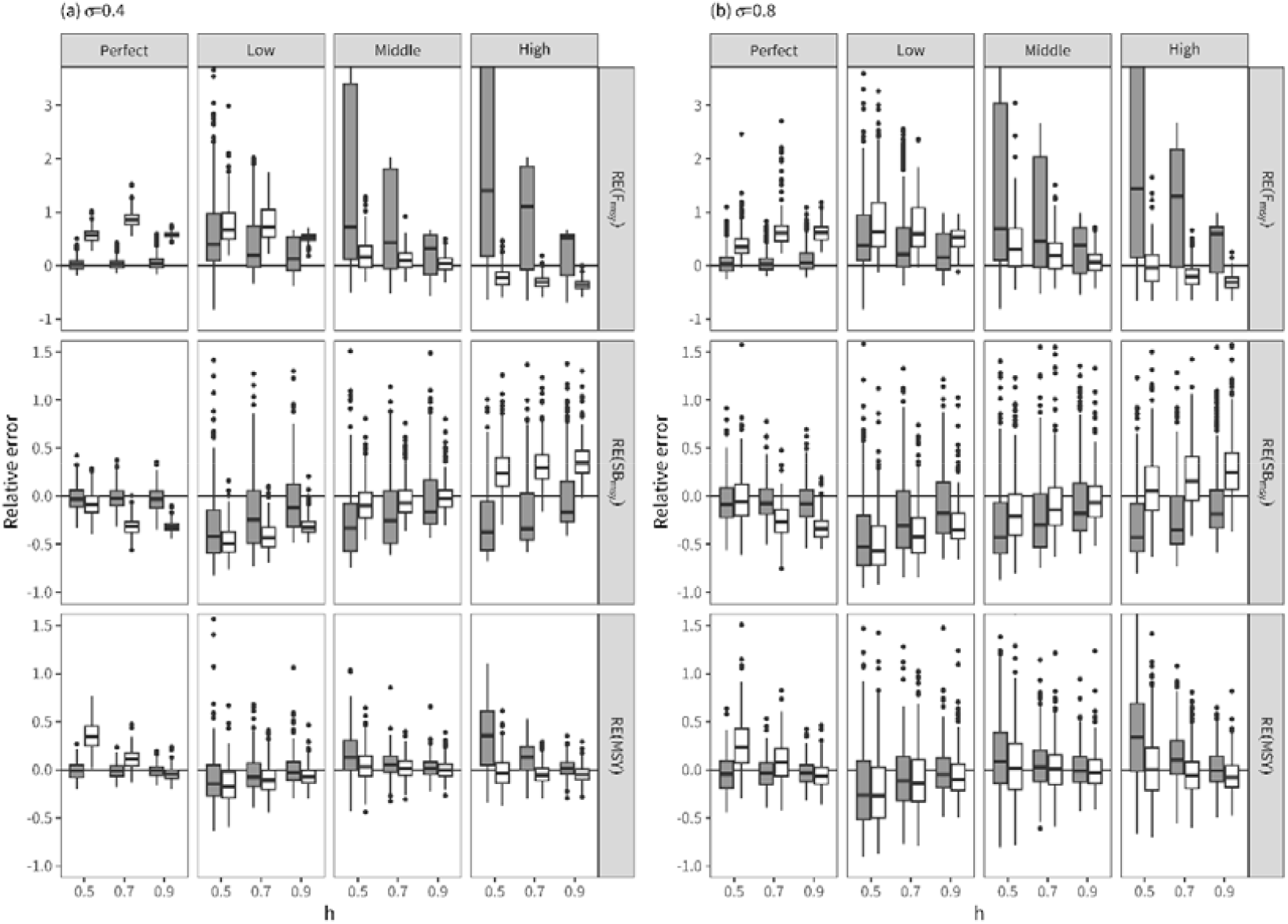
Relative errors of the MSY RP estimates of F_MSY_ (RE(F_MSY_)), SB_MSY_ (RE(SB_MSY_)), and MSY (RE(MSY)) when BH (gray) or HS (white) was used as the SRR for estimating the MSY RPs in the first management year under different data contrast scenarios (perfect, low, middle, and high) for the 1_SP species. The true SRR was assumed to be BH with varying steepness (*h* = 0.5, 0.7, 0.9) and recruitment variability (*σ* = 0.4, 0.8).

The incorrect use of HS when the true SRR was BH resulted in more biased estimates for the RPs in all the scenarios (Fig. 3, Figs. C1–C3). However, a notable feature of HS was that the variances of the relative errors tended to be smaller than those obtained under BH, particularly in the no-data-contrast scenarios. The variances exhibited under HS remained smaller than those observed under BH even in the middle and high scenarios and in the 2_SP and 3_MD scenarios, where the overall variances obtained under BH were particularly large (Figs. C1 and C2). With respect to the estimation biases, HS either overestimated or underestimated F_MSY_ depending on the scenario. Overestimation occurred under the “low” scenario, underestimation occurred under the “high” scenario, and small biases were observed under the “middle” scenario. The variances of the relative errors induced under *σ* = 0.8 were greater than those induced under *σ* = 0.4, although the general tendencies exhibited by the median relative errors remained similar to those observed for *σ* = 0.4.

### Long-term performance achieved when using HS instead of BH

The median relative errors and relative management metrics obtained for five different MPs were compared across different data contrast scenarios when *h* = 0.7 and *σ* = 0.4 in 1_SP (Fig. 4). When the population was depleted (the “low” scenario, Fig. 4, left column), the estimation bias of the MSY RPs was the smallest in MP-BH/N, as expected, since this MP correctly specified the SRR. For the other four MPs, in which the incorrect assumption of the SRR led to biased estimates of the MSY RPs, the biases tended to decrease over the management period, and some MPs achieved stock recovery to SB_MSY_. In addition, the catches of all four MPs remained near the MSY throughout the entire management period. Among the four MPs, MP-AIC/C performed better than the others and attained performance similar to that of MP-BH/N. During the initial phase of management under MP-AIC, HS was predominantly selected because the AIC difference was less than 2. As the data accumulated, BH was selected more frequently, resulting in a gradual reduction in the bias. In the middle scenario (Fig. 4, middle column), the biases of the MSY RP estimates were generally small and decreased over the management period, with all the MPs maintaining stock biomasses around SB_MSY_ and catches near MSY. In the “high” scenario, the biases of the MSY RPs did not improve under the four MPs when HS was used, with the initially underestimated F_MSY_ resulting in stock levels that were consistently above SB_MSY_. In contrast, under MP-BH/N, the initial overestimation of F_MSY_ was corrected through management, allowing the stock to be maintained around SB_MSY_.

**Fig. 4.**
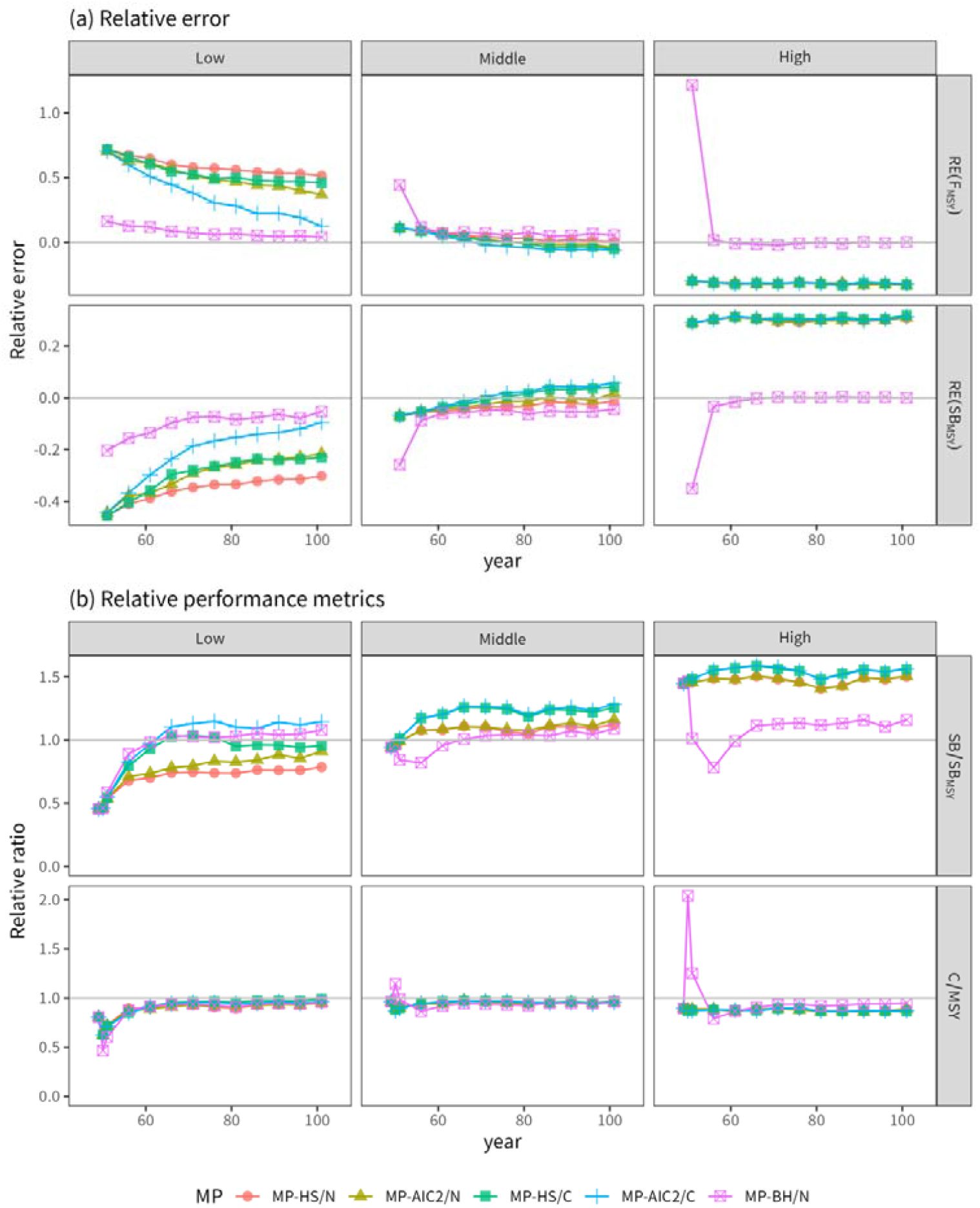
Median trajectories of (a) the relative errors RE(F_MSY_) and RE(SB_MSY_) and (b) the relative performance metrics of SB/SB_MSY_ and C/MSY when *h* = 0.7 and *σ* = 0.4 for the 1_SP species. The results obtained for management years 0–3 (years 49–51) and every 5-year interval when the MSY RPs were calculated are only shown for simplicity.

The degrees of bias improvement exhibited by the MSY RPs and the effectiveness of each MP varied with the species life history parameters. Figs. C4–C6 present the results obtained when the life history parameters of 2_MP, 3_MD, and 4_LD were used under the same *h* and *σ*, as shown in Fig. 4. While the relative performances of the five MPs remained consistent, the improvements in their bias and stock recovery results produced under the “low” scenario were slightly lower for 2_MP than for 1_SP and further reduced for 3_MD and 4_LD. In 3_MD and 4_LD, little difference was observed between MP-HS/C and MP-HS/N or between MP-AIC/C and MP-AIC/N, suggesting that capping was largely ineffective. MP-AIC still performed slightly better than MP-HS did, indicating a gradual increase in the correct BH selection rate over time. Nevertheless, gradual improvements continued across all the MPs, and simulations extended for more than 100 years revealed that the bias and stock recovery improvements continued under the “low scenario”, even in 3_MP and 4_LD (the associated results are not shown).

The comprehensive results obtained under all combinations of the tested parameters and prediction intervals (Fig. D) suggest that the speeds of bias correction and stock recovery were also dependent on the assumption of *h* and *σ* A lower *h* resulted in slower recovery, and greater recruitment variability made the MP with capping more effective.

The performance statistics calculated across all the parameters and species (Fig. 5) summarize the differences among the five MPs in terms of their initial catch reductions (Cinit), spawning biomass-related statistics (SBsafe, SB10, and SB30) and variability statistics (CAAV and SBmsyAAV), whereas the average catch differences (AC10 and AC30) were not significant. In terms of the biomass-related statistics, compared with the other three MPs, MP-HS/N and MP-AIC/N (without capping) performed relatively poorly, whereas MP-HS/C and MP-AIC/C (with capping) performed comparably to MP-BH/N in terms of the median. However, these two MPs showed greater variance, especially in SB40, than MP-BH/N did. The variance in SB40 reflects differences in recovery speed. As shown in Fig. D, recovery speed is influenced by life history traits and SR parameters; lower steepness, lower recruitment variability, and longer generation times tend to slow stock recovery (i.e. result in lower SB40). In terms of the variability statistics, compared with the other four MPs, MP-BH/N performed worse. The order of performance for CAAV, from best to worst, was MP-AIC/C, MP-HS/C, MP-AIC/N, MP-HS/N, and MP-BH/N. The performances achieved for SBmsyAAV among the four MPs other than MP-BH/N were broadly similar. In terms of the initial catch, compared with the other MPs, MP-BH/N yielded slightly lower catches and exhibited greater variance, reflecting greater variances in its estimates of F_MSY_.

**Fig. 5.**
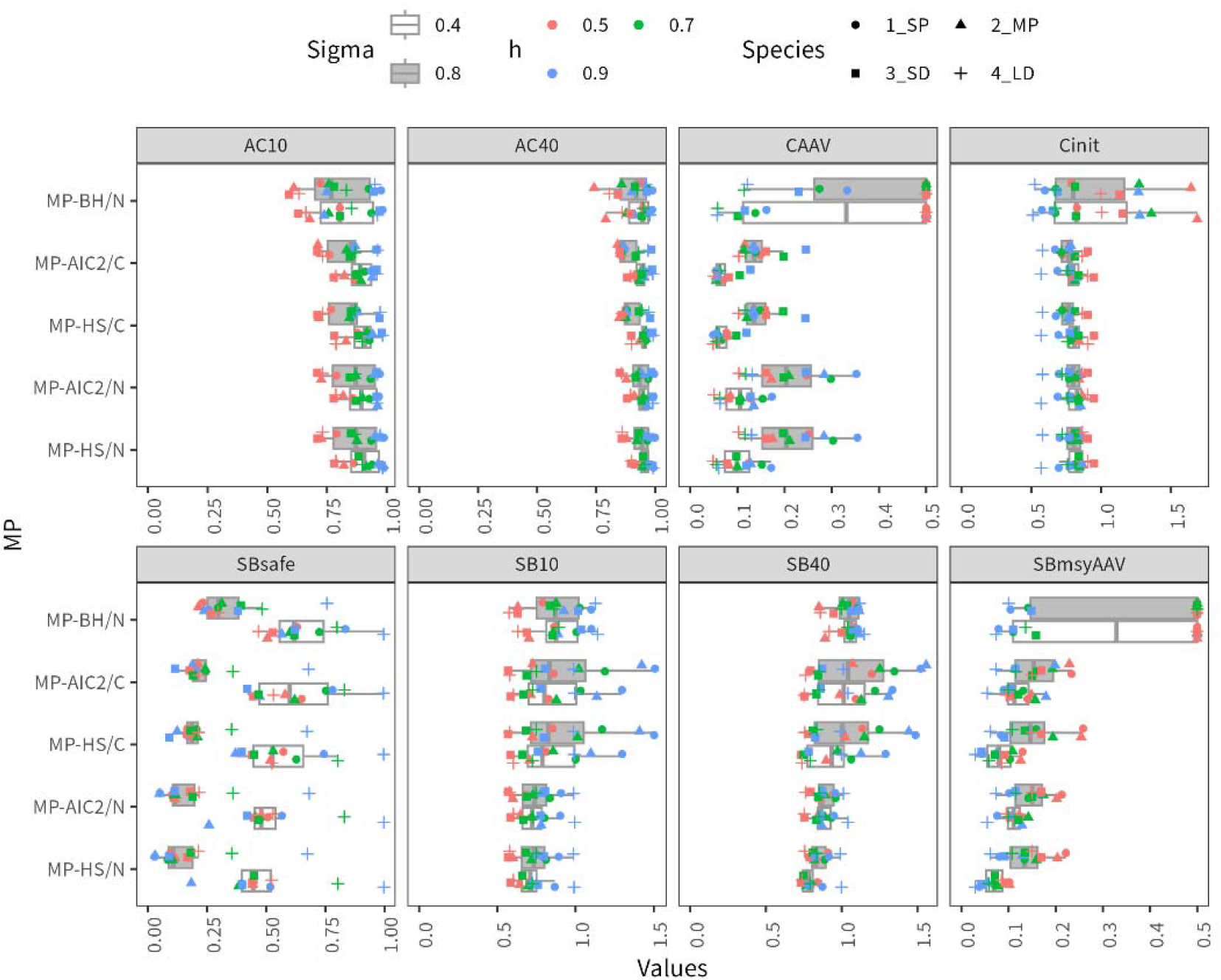
Comparison among eight performance statistics yielded by five MPs under the “low” scenario for all examined parameters and species. The maximum value of the y-axis for SBmsyAAV was set to 5; if it exceeded this value, it was plotted as 5.

The performances attained under the “middle” and “low” scenarios (Figs. C7 and C8, respectively) were generally consistent with the results shown in Fig. 5. Higher and more stable spawning biomass levels were achieved by MP-HS and MP-AIC than by MP-BH, as the underestimation of F_MSY_ by MP-HS and MP-AIC unintentionally led to more conservative management. The relative performance achieved in terms of the variability metrics (CAAV and SBmsyAAV) was also similar to that observed under the “low” scenario, with MP-BH showing greater variability than MP-HS or MP-AIC did. The average catch differences were not significant across the different MPs.

The estimated coefficients derived from the GLM analysis quantified the effects of the SR parameters, species, and management strategies on management success (Fig. 6). The effects of the SR parameters (*σ* and *h*) were consistent across the various stock depletion scenarios and species. The negative coefficients obtained with *σ* = 0.8 suggest that higher recruitment variability resulted in lower catch and spawning biomass. The negative and positive coefficients obtained with *h* = 0.5 and *h* = 0.9, respectively, suggest that higher *h* values resulted in higher catch and SSB.

**Fig. 6.**
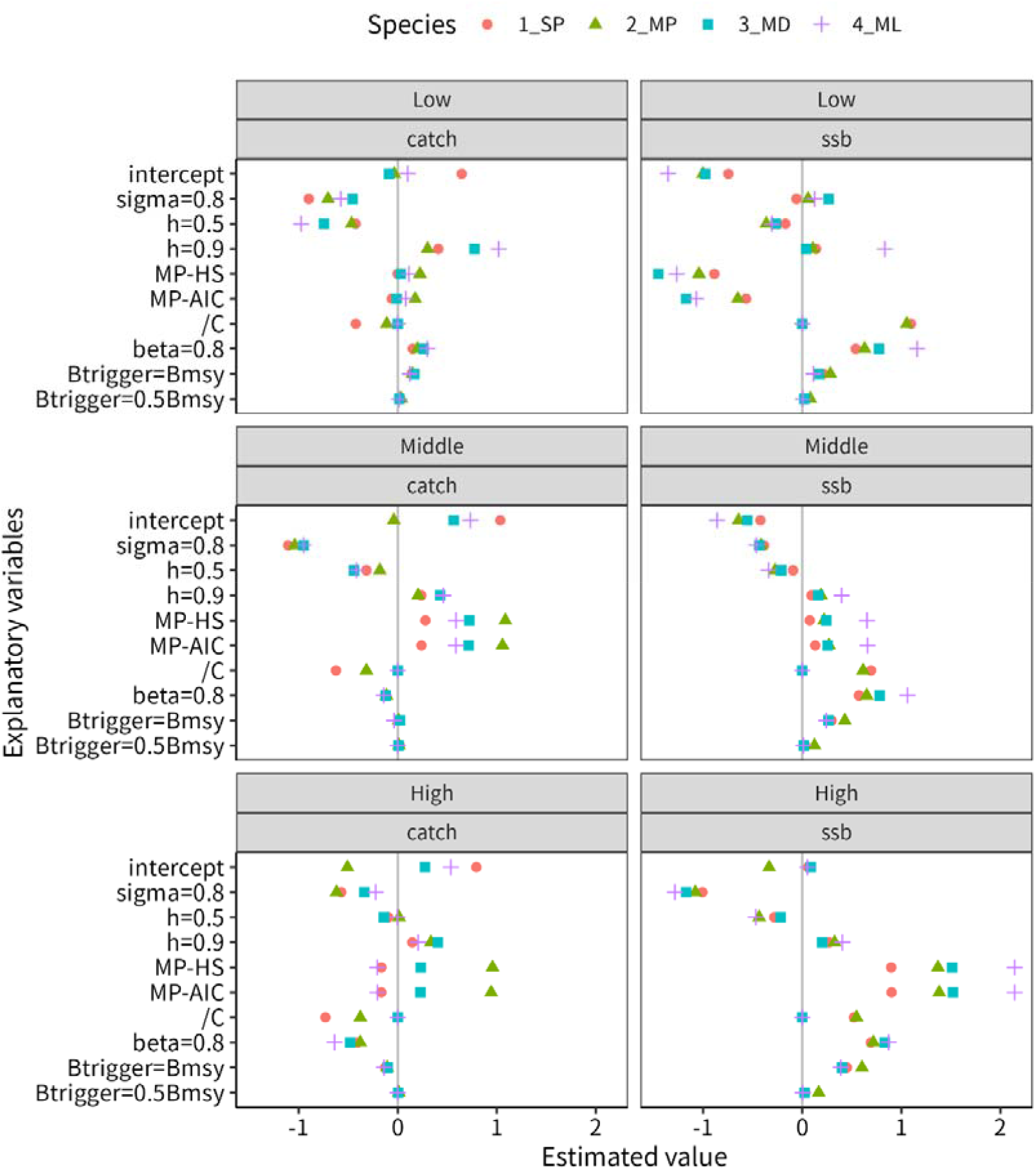
Estimated coefficients derived from the GLM analysis.

With respect to the effects of the management strategies under the “low” scenario, the negative coefficients of the strategies based on HS and AIC (MP-HS and MP-AIC, respectively) indicate reductions in the SB_success_ probability under these strategies (Fig. 6). For example, a coefficient of -1 corresponded to a decrease in probability from 50% to approximately 27%. In contrast, these strategies had little effect on Catch_success_, as their coefficients were close to zero. The effect of catch capping (/C) on SB_success_ depended on the species, with substantial positive effects observed for the pelagic species (1_SP and 2_MP) but little or no effect for the demersal species (3_MD and 4_ML). The increased probability of SB_success_ was associated with a reduction in Catch_success_, particularly in 1_SP, whereas this trade-off was less evident in the other species. Adopting a precautionary value of 0.8 (beta=0.8) increased SB_success_ and slightly increased Catch_success_ across all the species. In contrast, the effects of Btrigger (Btrigger=Bmsy and Btrigger=0.5Bmsy) were minor compared with those of the other strategies. For the “middle” and “high” scenarios, all the strategies except MP-HS and MP-AIC had effects that were consistent with those observed in the “low” scenario, whereas MP-HS and MP-AIC had opposite effects. Results of the sensitivity analysis assuming RI as the true SR relationship were broadly consistent with those obtained under the assumption that BH was true SR relationship (Appendix Fig. C9).

## Discussion

Our results show that when the true SRR follows the HS function, the stochastic MSY RPs are strongly influenced by recruitment variability, and the ratio of SB_MSY_ to SB_0_ spans a wider range than those derived from BH and RI. When HS is applied to a population that follows the BH SRR, SB_MSY_ tends to be underestimated, and F_MSY_ overestimated when the stock is heavily depleted and the data exhibit limited contrast. At the same time, MSY RPs derived from HS exhibit lower variance than those from BH, highlighting a bias–variance trade-off among alternative SRRs. In addition, the estimation biases in HS can be mitigated through adaptive management in which MSY RPs are periodically updated, allowing catches to remain near MSY over time. Importantly, effective mitigation within a realistic management timeframe requires adaptive management to be combined with precautionary strategies, such as applying an HCR with a precautionary factor of 0.8 and capping the amount of annual catches at the estimated MSY. Notably, compared with strategies based on the true BH SRR, HS-based strategies may offer the additional advantage of reducing catch variability and avoiding substantial initial catch reductions.

### Characteristics of MSY PRs under HS

Deterministic MSY RPs based on the HS SRR can be analytically derived from the relationships between SB_HS_, yield per recruit (YPR), and R_0_ (Mesnil et al. 2010, Okamura et al. 2021). Under deterministic population dynamics, if SB_MAX_ (the multiplier between R_0_ and SPR corresponding to the maximum YPR) is greater than SB_HS_, SB_MAX_ and F_MAX_ define SB_MSY_ and F_MSY_, respectively. Conversely, if SB_MAX_ is lower than SB_HS_, then SB_HS_ and the fishing mortality corresponding to SB_HS_ (F_HS_) become equivalent to SB_MSY_ and F_MSY_, respectively. Therefore, the upper bound of F_MSY_ under HS is set by F_MAX_. This theoretical framework was confirmed by our results obtained under *σ* = 0 (Fig. 2), which shows that the diagonal lines corresponding to SB_HS_ and the horizontal lines corresponding to SB_MAX_ jointly determine the position of SB_MSY_.

This study further examined the effects of recruitment stochasticity on MSY RPs under HS by defining stochastic MSY as the maximum long-term average yield obtained under population dynamics incorporating recruitment variability. The results showed that stochastic MSY RPs under HS were influenced by the level of *σ* (Fig. 2), whereas those under BH and RI were insensitive to *σ* due to the bias correction term (− 0.5 *σ*^2^) applied to annual recruitment deviations 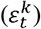 (Appendix A). Although the same bias correction term was applied under HS, HS exhibits the sensitivity to *σ* because recruitment variability can cause SB to fall below SB_HS_ even when fishing mortality is less than F_HS_. Once the biomass decreases below SB_HS_, recovery is delayed because of the absence of a compensatory effect below SB_HS_, resulting in reduced yield during the recovery period. Consequently, SB_MSY_ is estimated to be a higher level than SB_HS_

An exception occurs when *h* is very low. In such cases, maintaining SB above SB_HS_ requires substantially higher biomass levels, and the associated reduction in fishing mortality can result in greater yield loss than that induced by allowing the biomass to temporarily fall below SB_HS_. As a result, SB_MSY_ may be estimated below SB_HS_. Theoretically, there is no equilibrium below SB_HS_ under HS. However, the assumption of 10 times the generation time as equilibrium in this study allowed for transient states below SB_HS_. Such conditions are unlikely to occur in real population dynamics, and caution is needed when SB_MSY_ is estimated to be below SB_HS_.

The incorporation of recruitment stochasticity in calculating MSY RPs resulted in SB_MSY_ values that were, on average, 1.13 and 1.35 times higher SB_MSY_ under *σ* = 0.4 and 0.8, respectively, than those under deterministic conditions. This magnitude is comparable to the relationship between a biomass limit reference point (Blim, generally defined as SB_HS_ when HS is used) and a precautionary reference point (Bpa, generally defined as 1.39Blim by default) in ICES (Silvar-Valadomiu et al. 2021). This consistency suggests that stochastic MSY RPs under HS provide precautionary biomass levels that are similar to those adopted in the current ICES management frameworks. Furthermore, ICES does not apply MSY RPs derived from HS in a naïve manner but instead adopts a hierarchical approach that accounts for SRR uncertainty, thereby reducing the risk of bias arising from the use of any single SRR (ICES 2022). In Japan, stochastic MSY RPs under recruitment variability are adopted to calculate MSY RPs (FRA 2025b). A limitation of stochastic MSY RPs is their computational cost. However, Okamura et al. (2025) proposed an analytical method for calculating stochastic MSY RPs, which may facilitate their more practical and widespread use in the future.

A target biomass level of 30–40% of SB_0_ is typically used as a proxy for SB_MSY_ when direct estimation is difficult. Our results under BH and RI support the validity of this range. In contrast, SB_MSY_/SB_0_ under HS spanned a much wider range and frequently exceeded 0.5 when *h* was less than approximately 0.5 (Fig. 2). These results are consistent with those of Punt et al (2014), who reported that the spawning biomass ratio achieving maximum economic yield (SB_MEY_) to SB_0_ was broader and generally greater under HS than under BH or RI. Because SB_MEY_ is typically higher than SB_MSY_, the broader distribution of SB_MEY_ suggested by Punt et al. (2014) likely reflects the wider range of SB_MSY_ under HS. These results indicate that the commonly used proxy of 30–40% of SB_0_ may not be applicable when using HS. In contrast, %SPR_MSY_ was similar across the three SRRs, suggesting that %SPR_MSY_ is less sensitive to the choice of SRR, including HS. These results highlight the notion that the empirical proxy RPs established under BH or RI cannot be applied without careful evaluation when HS is used as the SRR.

The observed ranges of SB_MSY_/SB_0_ and %SPR_MSY_ depend on the assumed steepness range *h*) and should therefore be interpreted with caution. This is particularly important for HS, where the definition of *h* differs from those of BH and RI. Under HS, *h* < 0.5 implies that there is no recruitment compensation below half of SB_0_, which may be biologically unrealistic. Empirical understanding of plausible *h* for cases when the HS function is applied needs to be obtained. Similarly, for RI, *h* values greater than one were examined to explore a wide range of possible dynamics, but the biological plausibility of such values remains uncertain. Because most empirical meta-analyses involving *h* have been conducted under the assumption of the BH relationship (e.g., Thorson 2020), further empirical work is needed to evaluate *h* and its biological interpretation under alternative stock–recruitment relationships, including HS and RI.

### Robustness of the HS SRR in operational use for management purposes

Our results show that, under insufficient data contrast, the HS SRR is characterized by a large bias but low variance, whereas the BH SRR exhibits a smaller bias but higher variance (Fig. 4). This pattern reflects a bias–variance trade-off among alternative SRRs. This trade-off arises from the structural property of HS that avoids extrapolating beyond the observed range of recruitment, thereby preventing unrealistic management reference point estimates. While HS has been proposed as a pragmatic alternative to conventional SRRs (e.g., Clark et al. 1985; Barrowman and Myers 2000; Mesnil et al. 2010; Ichinokawa et al. 2017), our results provide simulation-based support for its use, showing that avoiding extrapolation reduces variance in MSY-based reference point estimates, albeit at the cost of increased bias under insufficient data contrast.

A key consideration when the HS SRR is applied in fishery management is that using the HS SRR under insufficient data contrast can lead to biased MSY RP estimates. This underscores the importance of adopting precautionary and adaptive management when applying HS SRR. Our results (Fig. 6) show that relying solely on HS does not ensure convergence to the true MSY state when the true SRR follows BH, particularly under heavily depleted conditions, even as data accumulate over time. This bias arises primarily from the persistent overestimation of F_MSY_, despite partial stock recovery and yields approaching the true MSY. However, precautionary measures, such as applying a precautionary factor of 0.8 to the HCR and capping annual catches at the estimated MSY, combined with adaptive model updating, can mitigate these biases and enhance the stock recovery (Figs. 5–6). The bias reduction rate depends on the life history parameters, *h*, and *σ*. Notably, the adaptive transition from an initial HS assumption towards the true BH SRR outperformed the strategy of assuming BH from the outset, particularly in terms of the stability of the resulting MSY estimates and the magnitudes of the initial catch reductions (Fig. 5). In addition, catch capping alone contributed significantly to stabilizing the annual catches, supporting its effectiveness as a robust precautionary tool.

However, as shown in Figs. 5–6, the direction and magnitude of the bias in the MSY RP estimates under HS strongly depend on stock status. In particular, when the stock is well above SB_MSY_, the MSY RPs estimated under HS may become overly conservative. It is informative to compare MSY RPs derived across different SRRs with other commonly used biological reference points, such as %SPR. The upper bound of F_MSY_ under HS, imposed by F_MAX_, can limit excessive fishing mortality. However, because F_MAX_ depends on selectivity and tends to increase when selectivity favours older ages (Wang et al. 2014), excessively large F_MAX_ values may weaken this safeguard. A similar issue may arise under BH when *h* is close to 1.

In this study, three precautionary management strategies, i.e., catch capping, a precautionary factor of 0.8, and Btrigger, were compared, and the results demonstrated that their effectiveness strongly depended on the life history characteristics of the examined species (Fig. 6). Catch capping was particularly effective for pelagic species, substantially increasing the probability of management success, whereas its effect was minimal for demersal species. This difference likely reflects generation time differences. In pelagic species with short generation times, recruitment fluctuations were rapidly transmitted to the total biomass and, consequently, to TAC. Under such conditions, TAC may frequently exceed the estimates of MSY, and catch capping effectively reduces the catch levels, thereby promoting stock recovery. In contrast, for demersal species with longer generation times and more buffered population dynamics, TAC rarely exceeded the estimates of MSY even without catch capping, limiting its additional benefit.

The precautionary factor (β = 0.8) was particularly effective for demersal species and provided consistent benefits for pelagic species, suggesting that directly reducing the realized fishing mortality rate is a broadly robust precautionary approach. In Japan, the use of a precautionary adjustment factor of β = 0.8 has been recommended as a standard component of HCR (Okamura et al. 2020, FRA 2025c). In contrast, adjusting B_trigger_ had relatively limited effects on achieving MSY-based reference points although B_trigger_ might remain important for preventing extreme stock depletion by reducing the fishing mortality rate at low biomass levels. Our results suggest that precautionary factors are broadly effective across species, whereas catch capping may be particularly beneficial for species with high recruitment variability and rapid population turnover. More generally, the choice of precautionary strategy should consider the life history characteristics of species and management objectives.

This simulation study simplifies the complexity of real-world fisheries stocks and management dynamics, as its primary goal is to illustrate the potential utility of using the HS SRR as an expedient assumption in combination with adaptive learning and precautionary strategies. While MSE frameworks commonly incorporate sources of uncertainty, including stock assessment errors, implementation errors, and autocorrelations (Punt et al. 2016), these factors were not incorporated into this study to isolate the minimum possible bias arising from limited data contrast and model misspecifications. As such, the precautionary management strategies proposed in this study should be regarded as minimum baselines under realistic fisheries stocks with uncertainty.

The concept of MSY as a management reference point experienced renewed interest in the early 2000s (Mace 2001). Several fisheries management organizations that had previously been cautious about MSY began adopting MSY-based approaches (Ichinokawa et al. 2017, Mesnil 2012, Okamura 2023). The accumulation of best practices in MSY-based management (Hilborn and Ovando 2014; Worm et al. 2009) has facilitated its broader adoption, including in regions where stock assessment and fisheries management remain under development (Costello et al. 2012). This study supports this broader transition by demonstrating that the pragmatic use of HS, when combined with a precautionary strategy, can serve as a practical entry point for implementing MSY-based management in fisheries where unstable MSY estimates have hindered its application. The HS-based approach can also reduce the magnitude of the initial catch reduction and the variability of the estimated MSY RPs, thereby potentially improving stakeholder acceptance. However, it should also be noted that the use of HS is more effective when it is accompanied by precautionary strategies, such as the application of precautionary adjustment factors and catch caps, for mitigating potential biases. These findings highlight the value of combining the pragmatic HS SRR with precautionary and adaptive management for supporting MSY-based fisheries management under uncertainty.

## Use of AI tools

The authors used AI-assisted tools (Microsoft M365 Copilot) to support the English proofreading and language refinement processes during the manuscript preparation phase. All scientific content, analyses, interpretations, and conclusions were developed entirely by the authors, and the authors take full responsibility for the final manuscript.

## Author contributions

Momoko Ichinokawa was responsible for developing and implementing all the simulation codes, conducting the analyses, organizing and summarizing the results, and drafting the initial version of the manuscript. Hiroshi Okamura worked together with Ichinokawa to formulate the overall research direction, conceptual design, and interpretation of the findings and contributed to refining the structure of the manuscript. Both authors discussed the methodological approach, shaped the arguments presented in the paper, and approved the final submitted version.

