## Appendix for "Robustness and management performance of MSY reference points derived from the hockey-stick stock-recruitment model under structural uncertainty"

Running head: HS-derived MSY reference points

Key word: stock-recruitment relationship, hockey-stick, segmented regression, management strategy evaluation, maximum sustainable yield, precautionary approach, bias variance trade-off,

1. Fisheries Resources Institute, Japan Fisheries Research and Education Agency, 2-12-4 Fukuura, Kanazawa-ku, Yokohama, Kanagawa 236-8648 Japan
2. School of Data Science, Yokohama City University, 22-2 Seto, Kanazawa-ku, Yokohama, Kanagawa 236-0027 Japan

### Appendix

#### Appendix A. Equations for the population dynamics of the operating models and defining the MSY reference points

##### List of symbols

| | |
|---|---|
| $y$ | Year $y$ |
| $a$ | Age, $a = A_{\min}, \ldots, A_{max}$ |
| $k$ | Iteration of the simulation |
| $N_{y,a}^{k}$ | Number of fish at year y, age $a$ and iteration $k$ |
| $F_{y}^{k}$ | Maximum fishing mortality coefficient at year $y$ and iteration $k$ |
| $M_a$ | Natural mortality coefficient at age $a$ |
| $m_a$ | Maturity rate at age $a$ |
| $w_a$ | Weight at age $a$ |
| $G_a$ | Fishing selectivity rate at age $a$ |
| $SB_{y}^{k}$ | Spawning biomass at year $y$ and iteration $k$, defined as $\sum_{a=A_{\min}}^{A_{max}} N_{y,a} w_{y,a} m_{y,a}$ |
| $C_{y}^{k}$ | Total catch weight at year $y$ and iteration $k$ |
| $\varepsilon_{y}^{k}$ | Recruitment deviations at year $y$, defined as : $\varepsilon_y \sim \mathrm{N}(-0.5\sigma^2, \sigma^2)$ |
| $\alpha_1, \alpha_2$ | Parameters in the SR function |

##### Population dynamics

The population dynamics assumed in the simulation model were structured by age. Recruitment occurred at age $a = A_{\min}$ according to the assumed SR function $f$ and the spawning biomass $SB_{t}^{k}$ as follows:

$$N_{y,A_{\min}}^{k} = f\left(SB_{y}^{k}, \alpha_1, \alpha_2\right) \exp\left(\varepsilon_{t}^{k}\right) \quad s = 0 \qquad \text{(A1).}$$

The annual recruitment deviation $\varepsilon_{y}^{k}$ was randomly generated from $\mathrm{N}(-0.5\sigma^2, \sigma^2)$, the parameters $\alpha_1$ and $\alpha_2$ are parameters of the SR function, and $\sigma$ is the standard deviation of the recruitment variability. The spawning biomass $SB_{t}^{k}$ was defined as $\sum_{a=A_{\min}}^{A_{\max}} N_{y,a}^{k} w_a m_a$, where $w_a$ and $m_a$ are the weight and maturity at age $a$, respectively. As the SR function $f$, the following three SR functions were considered: Hockey-Stick (HS, Clark et al. 1985, Eq. A2), Beverton–Holt (BH, Beverton and Holt 1957, Eq. A3), and Ricker (RI, Ricker 1954, Eq. A4), with 2 parameters.

$$N^k_{y,A_{\min}} = \begin{cases} \alpha_1 \times SB^k_{t-A_{\min}} & if\ SB^k_{y-A_{\min}} < b \\ \alpha_1 \times \alpha_2 & if\ SB^k_{y-A_{\min}} \geq b \end{cases} \quad (A2)$$

$$N_{y,A_{\min}} = \frac{a_1 SB^k_{y-A_{\min}}}{\left(1 + \alpha_2 SB^k_{y-A_{\min}}\right)} \quad (A3)$$

$$N_{y,A_{\min}} = a_1 SB^k_{y-A_{\min}} \exp\left(-\alpha_2 SB^k_{y-S_{\min}}\right) \quad (A4)$$

The parameters $\alpha_1, \alpha_2$ were determined from the assumed steepness $h$ and an average number of recruits without fishing ($R_0$), which was fixed to 100.

The dynamics of older fish ($a > A_{\min}$) were driven by natural mortality $M_a$, the fishing mortality coefficient $F^k_y$ and the selectivity rate $G_a$ as follows:

$$N^k_{y,a} = \begin{cases} N^k_{y-1,a-1} \exp\left(-M_{a-1} - F^k_{y-1} G_{a-1}\right) & A_{\min} < a < A_{\max} \\ N^k_{y-1,a-1} \exp\left(-M_{a-1} - F^k_{y-1} G_{a-1}\right) + N^k_{y,a} \exp\left(-M_a - F^k_y G_a\right) & a = A_{\max} \end{cases} \quad (A5).$$

The catch weights at age $C^k_{y,a}$ were calculated using the Baranov catch equation as follows:

$$C^k_{y,a} = \frac{F^k_y G_a}{M_a + F^k_y G_a} \left(1 - \exp\left(-F^k_y G_a - M_a\right)\right) N^k_{y,a} w_a \quad \text{(A6).}$$

#### Step A

We conducted 1000 simulations ($k = 1, \ldots, 1000$) with different random seeds using the above equations and all combinations of the biological parameters for 4 species, 3 types of SR functions, and scenarios involving steepness and recruitment variability to calculate the true MSY RPs under stochastic population dynamics. The simulations were carried out for 47, 55, 74, and 66 years for species 1–4, respectively, under the assumption that 10 times the generation time represents the equilibrium state. During this simulation, the fishing mortality coefficient $F^k_t$ was kept constant at $F^k_t = F$. Defining the final year of the simulations, i.e., $t = T_L$ , MSY was defined as $\max_{0<F} \sum_{k=1}^{1000} \sum_{a=A_{\min}}^{A_{\max}} C^k_{T_L,a} / 1000$, and $F_{MSY}$ was defined as $\operatorname{argmax}_{0<F} \sum_{k=1}^{1000} \sum_{s=A_{\min}}^{A_{\max}} C^k_{T_L,a} / 1000$ . Then, $SB_{MSY}$ is defined was the average spawning biomass when $F = F_{MSY}$.

#### Steps B and C

We categorized the simulation period as the burn-in period before the stock assessment ($T_0 < y < T_1$), the stock assessment period ($T_1 \leq y < T_2$), and the stock management period ($T_2 \leq y \leq T_3$). The fishing mortality coefficients $F_t^k$ before the stock management period ($y < T_2$) were determined according to the depletion scenarios, whereas those yielded during the stock management period ($T_2 \leq y < T_3$) were determined according to the harvest control rules in Step C.

The historical depletion levels of the stock were controlled by the parameter $P_y$: the relative ratio of the spawning biomass to $SB_{MSY}$ in year $y$ of the stock assessment period under deterministic population dynamics. We first defined the "perfect" scenarios with 100 years of observations, where $P_y = 0.05$, and the spawning biomass in the first year of the stock assessment was equal to $SB_0$. In the perfect scenario, fishing mortality was assumed to linearly increase from $y = 20$ to $y = 100$ to clarify the data contrast level. Then, realistic scenarios with no data contrast and 31 years of observations were considered, where the stock level remained constant at $P_{20} = P_{50}$ under constant fishing mortality. The stock levels at which the simulated population remained were $0.5B_{MSY}$ (low, $P_{20} = P_{50} = 0.5$), $1.0B_{MSY}$ (middle, $P_{20} = P_{50} = 1$), and $1.5B_{MSY}$ (high, $P_{20} = P_{50} = 1.5$) (Fig. A2).

In step B, we investigated the estimation biases of the MSY RPs in the first year of stock management $T_2$. During the actual procedure, a subset of the data derived from the observed recruitment $N_{y,A_{\min}}^{obs,k} = N_{y,A_{\min}}^{k}$ $(y = T_1, \ldots, T_2 - 1)$ and spawning biomass $SB_y^{obs,k} = SB_y^k$ $(y = T_1, \ldots, T_2 - 1)$ were applied to the SR function $f'$ to estimate the SR parameters $\widehat{\alpha_1}$ and $\widehat{\alpha_2}$ and calculate the stochastic MSY RPs with the estimated parameters by conducting the stochastic simulation described in Step A. Parameter estimation was conducted for the SR function by using the maximum likelihood method, i.e., maximizing the log-likelihood function of $(T_2 - T_1)\ln\left(\frac{1}{\sqrt{2\pi\varsigma}}\right) -$

$\frac{1}{2}\sum_{t=T_1}^{T_2-1}\left(\log\left(N_{y,A_{\min}}^{obs,k}\right) - \log\left(f'\left(SB_y^{obs,k},\alpha_1,\alpha_2\right)\right) + 0.5\varsigma^2\right)$. Then we calculated MSY RPs using the estimated SR parameters and definitions described in step A; however, the number of stochastic simulations was reduced to 500 to reduce computational time.

When the true and estimated MSY RPs were considered to be $\vartheta$ and $\hat{\vartheta}_y^k$, respectively, the relative bias of the estimated MSY RPs was calculated as $\frac{\hat{\vartheta}_y^k-\vartheta}{\vartheta}$. The true MSY RPs $\vartheta$ were considered those calculated in Step A. Here, the estimated reference points are represented as $\widehat{\mathrm{SB}}_{\mathrm{MSY}_y}^{\ k}$, $\widehat{\mathrm{F}}_{\mathrm{MSY}_y}^{\ k}$, $\widehat{\mathrm{MSY}}_y^k$, and $\widehat{\mathrm{SB}}_{0_y}^{\ k}$.

In Step C, the fishing mortality rate $F_y^k$ during the management period $(t \geq T_2)$ was determined from the harvest control rules and the estimated MSY RPs, which were updated every 5 years.

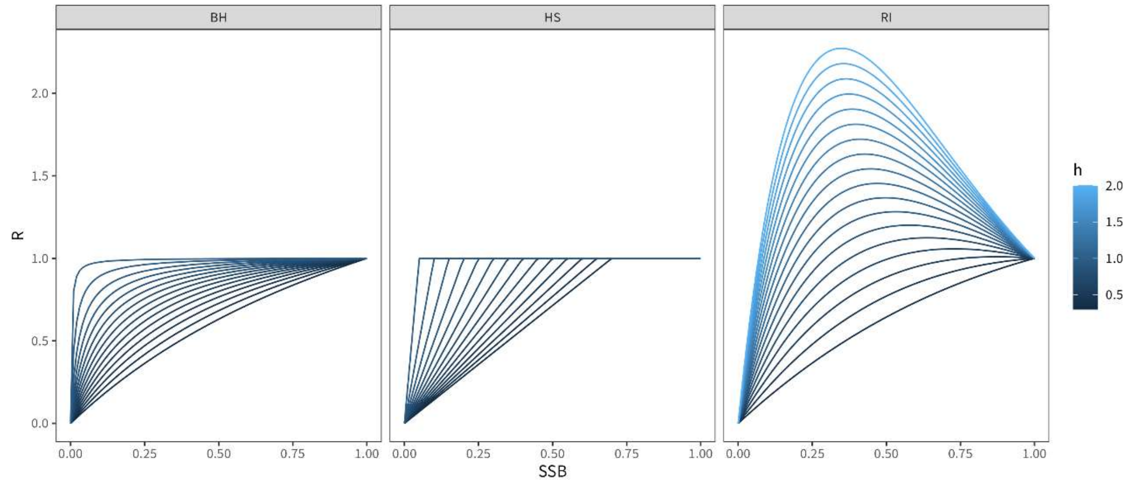


Fig. A1. Ranges of the variations exhibited by the SRRs assumed when calculating the MSY reference points.

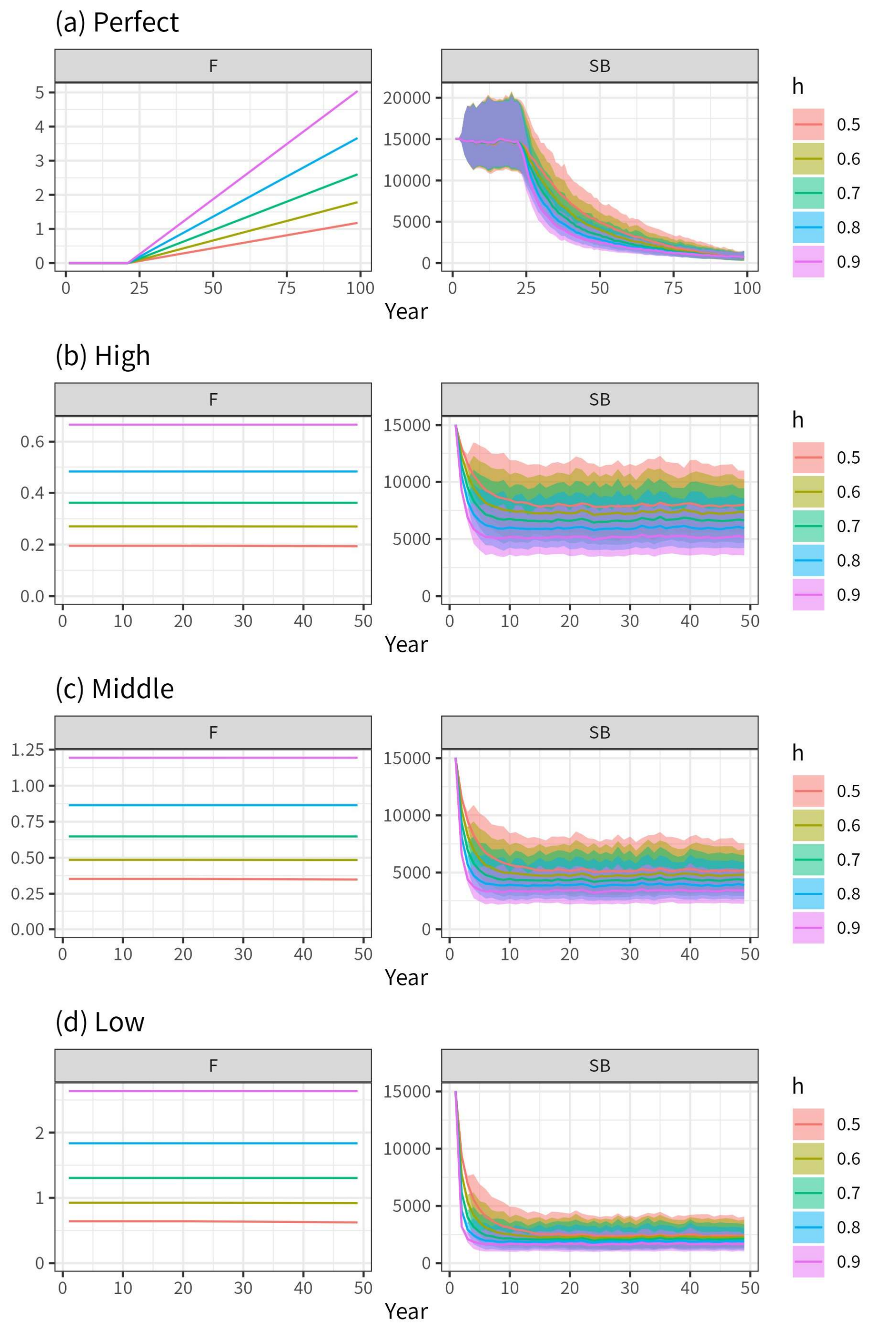


Fig. A2. Trajectories of the fishing mortality rate (left colums) and spawning biomass (right columns) under the “perfect”, “high”, “middle”, and “low” scenarios for species 1_SP and $\sigma^2$=0.4. In the “perfect” scenario, SRR data were collected from the 20th year to the 100th year, whereas in the other scenarios, the observation period ranged from the 20th year to the 50th year.

#### Appendix B. Biological parameters used in the simulation models

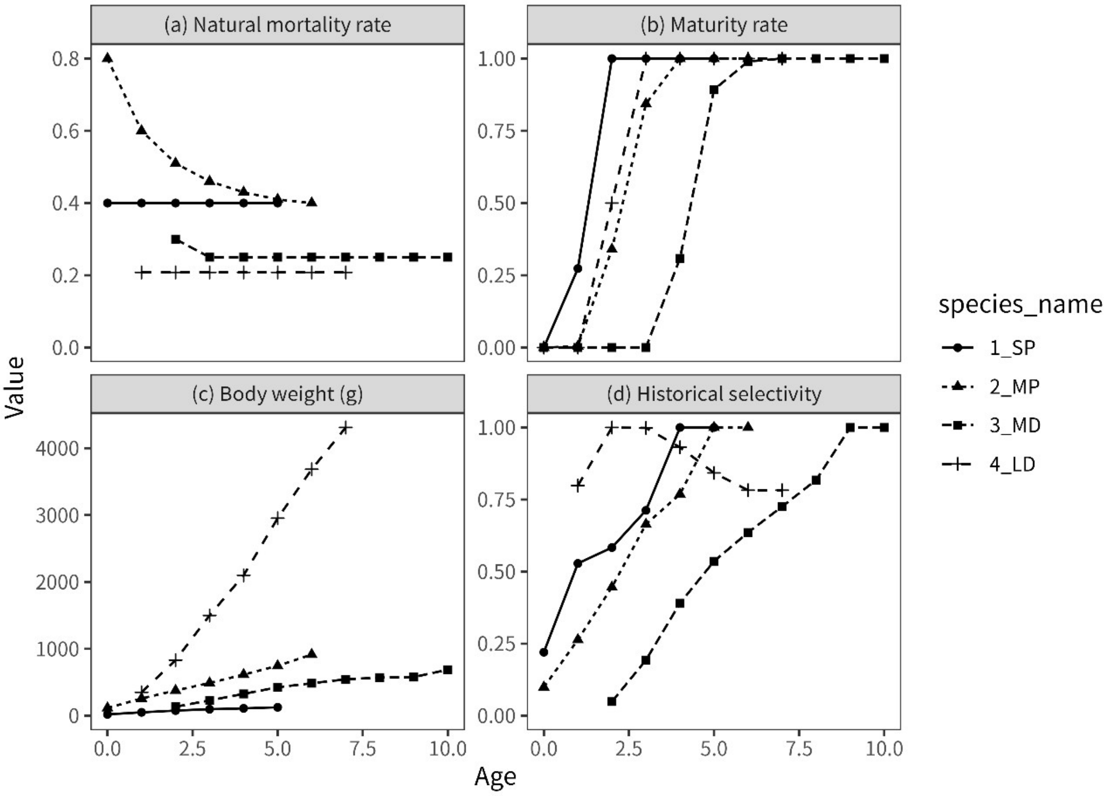


Fig. B. Biological parameters of the natural mortality rate (a), maturity rate (b), body weight (c), and historical selectivity rate (d) at age. 1_SP: Japanese sardine (small pelagic) (Furuichi et al. 2024), 2_MP: chub mackerel (medium pelagic) (Yukami et al. 2024), 3_MD: walleye pollock (medium demersal) (Chimura et al. 2024), and 4_LD: Japanese flounder (large demersal) (Masubuchi et al. 2024). Note that the fishery selectivity rate in the base case was assumed to be the same as the maturity rate, with a minimum of 0.1, and the historical selectivity rate shown in (d) was used for sensitivity analysis purposes. Some biological parameters and all selectivity rates were time-varying, but we used averaged values across all the years.

Furuichi, S., Yukami, R., Higashiguchi, K., Kamimura, Y., Nishijima, S., Watabe, R., & Isu, S. (2024). *Stock assessment and evaluation for the Pacific stock of Japanese sardine (2023).*

*Marine fisheries stock assessment and evaluation for Japanese waters. National Research Institute of Fisheries and Education Agency, Tokyo, Japan.* https://abchan.fra.go.jp/wpt/wp-content/uploads/2025/03/details_2024_01.p

Yukami, R., Nishijima, S., Kamimura, Y., Isu, S., Furuichi, S., Watabe, R., Higashiguchi, K., Saito, R., Ishikawa, K. (2024). *Stock assessment and evaluation for the Pacific stock of chub mackerel (2023). Marine fisheries stock assessment and evaluation for Japanese waters. National Research Institute of Fisheries and Education Agency, Tokyo, Japan.* https://abchan.fra.go.jp/wpt/wp-content/uploads/2025/03/details_2024_05.pdf

Chimura, M., Chiba, S., Sakai, O., Hamabe, K., Sato, R., and Hamazu, Y. (2024). *Stock assessment and evaluation for the Sea of Japan stock of walleye pollock (2023). Marine fisheries stock assessment and evaluation for Japanese waters. National Research Institute of Fisheries and Education Agency, Tokyo, Japan.* https://abchan.fra.go.jp/wpt/wp-content/uploads/2025/03/details_2024_09.pdf

Masubuchi, T., Iseki, T., Sakai, T., & Gomi, S. (2024). *Stock assessment and evaluation for the western central Sea of Japan and Tsushima current stock of bastard halibut (2023). Marine fisheries stock assessment and evaluation for Japanese waters. National Research Institute of Fisheries and Education Agency, Tokyo, Japan.* https://abchan.fra.go.jp/wpt/wp-content/uploads/2025/03/details_2024_63.pdf

### Appendix C. Figures demonstrating the results

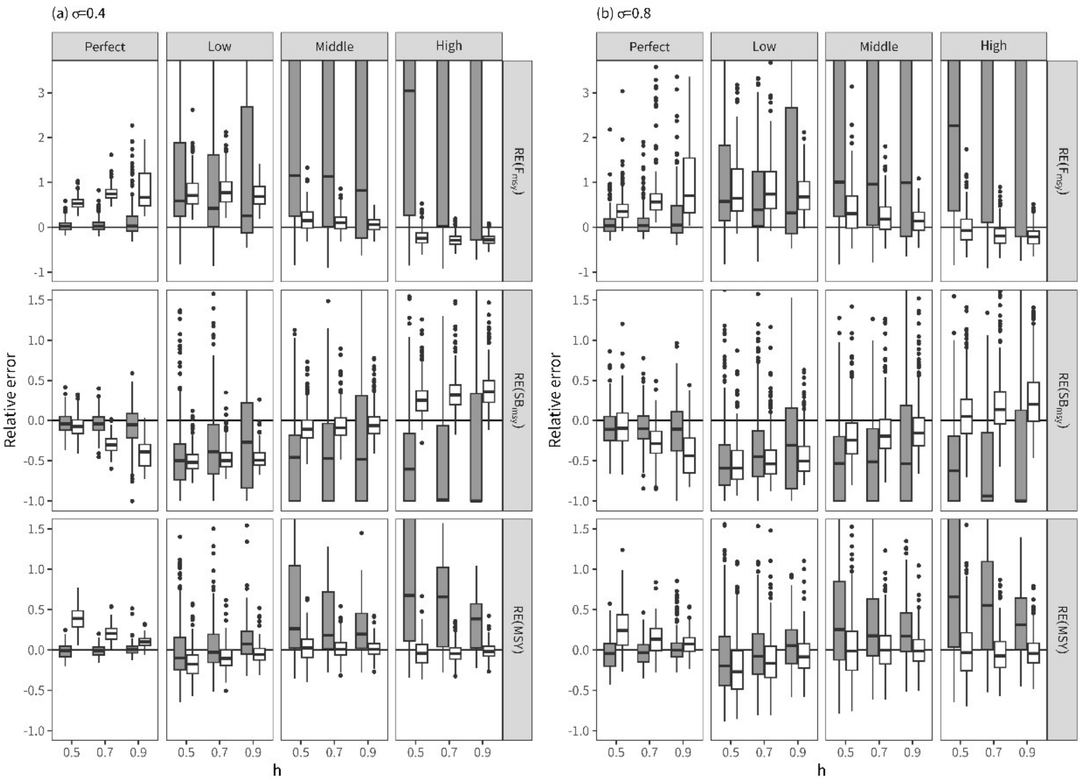


Fig. C1. Relative errors of the MSY RP estimates of $F_{MSY}$ (RE($F_{MSY}$)), $SB_{MSY}$ (RE($SB_{MSY}$)), and MSY (RE(MSY)) when BH (gray) or HS (white) was used as the SRR for estimating the MSY RPs in the first management year under different data contrast scenarios (perfect, low, middle, and high) for the 2_MP species. The true SRR was assumed to be BH with varying steepness ($h$ = 0.5,0.7,0.9) and recruitment variability (σ = 0.4,0.8).

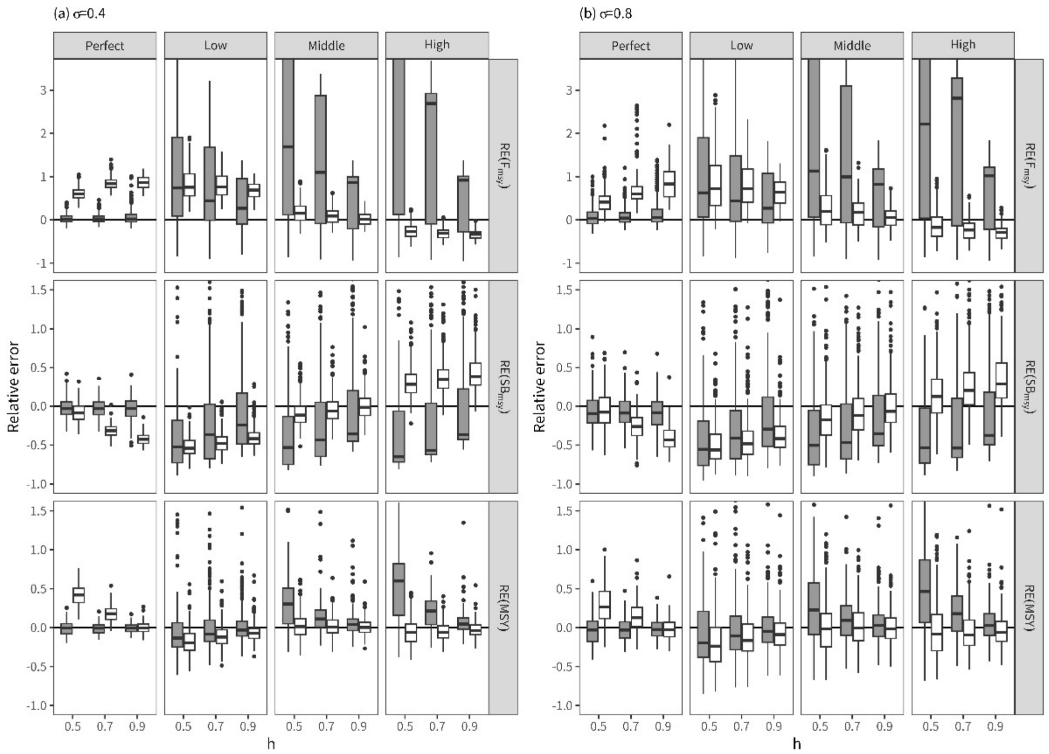


Fig. C2. Same as Fig. C1, but for the 3_MD species.

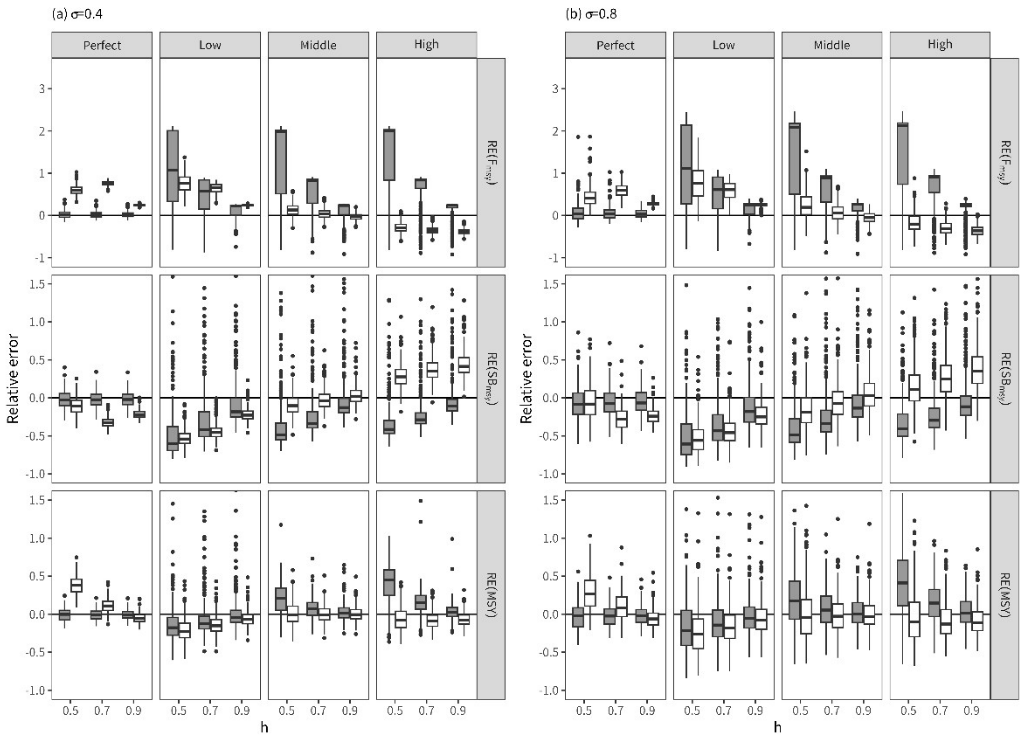


Fig. C3. Same as Fig. C1, but for the 4_MD species.

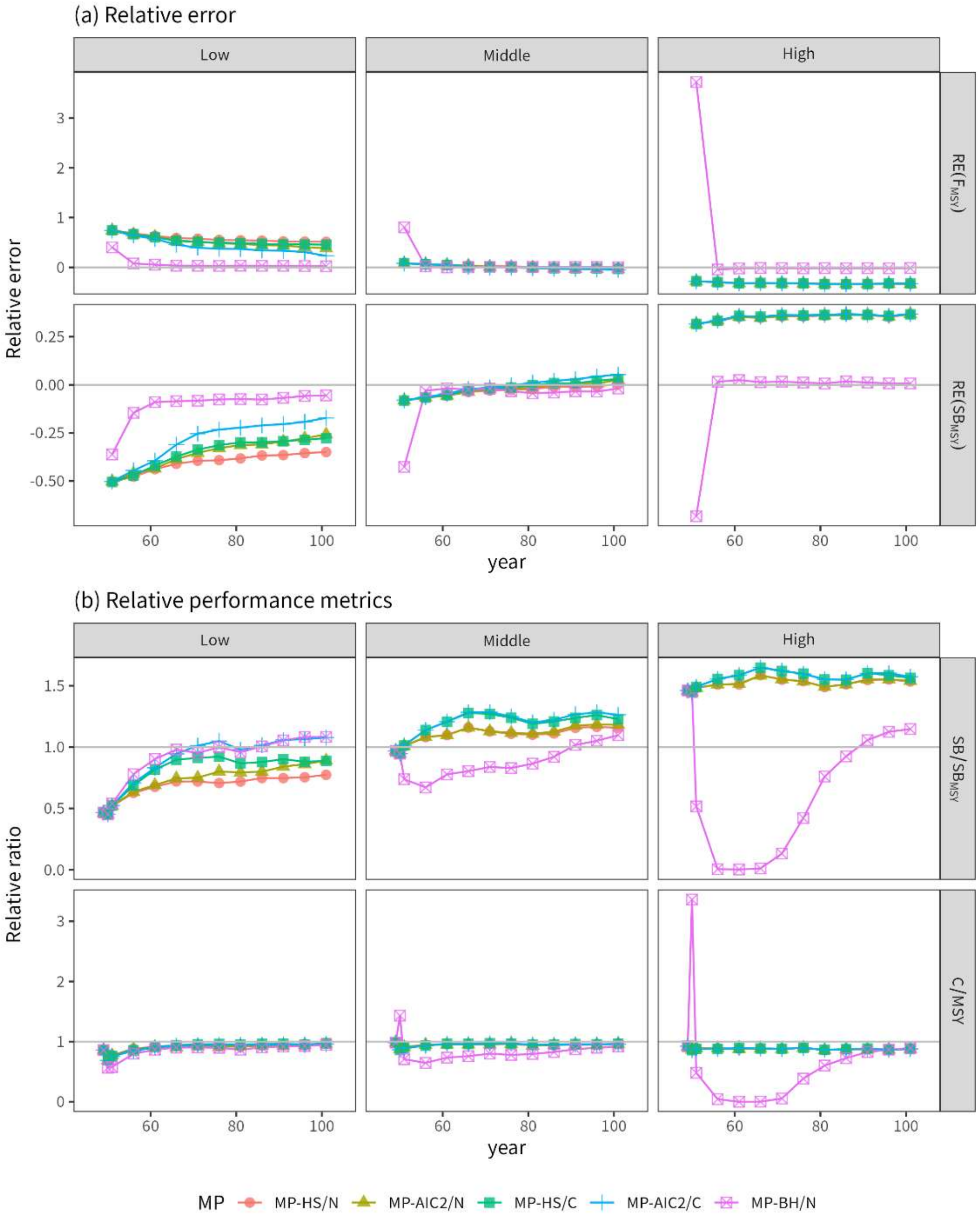


Fig. C4. Median trajectories of (a) the relative errors RE(FMSY) and RE(SBMSY) and (b) the relative performance metrics of SB/SBMSY and C/MSY when h=0.7 and σ=0.4 for the 2_MP species. The results obtained for management years 0–3 (years 49–51) and every 5-year interval when the MSY RPs were calculated are only shown for simplicity.

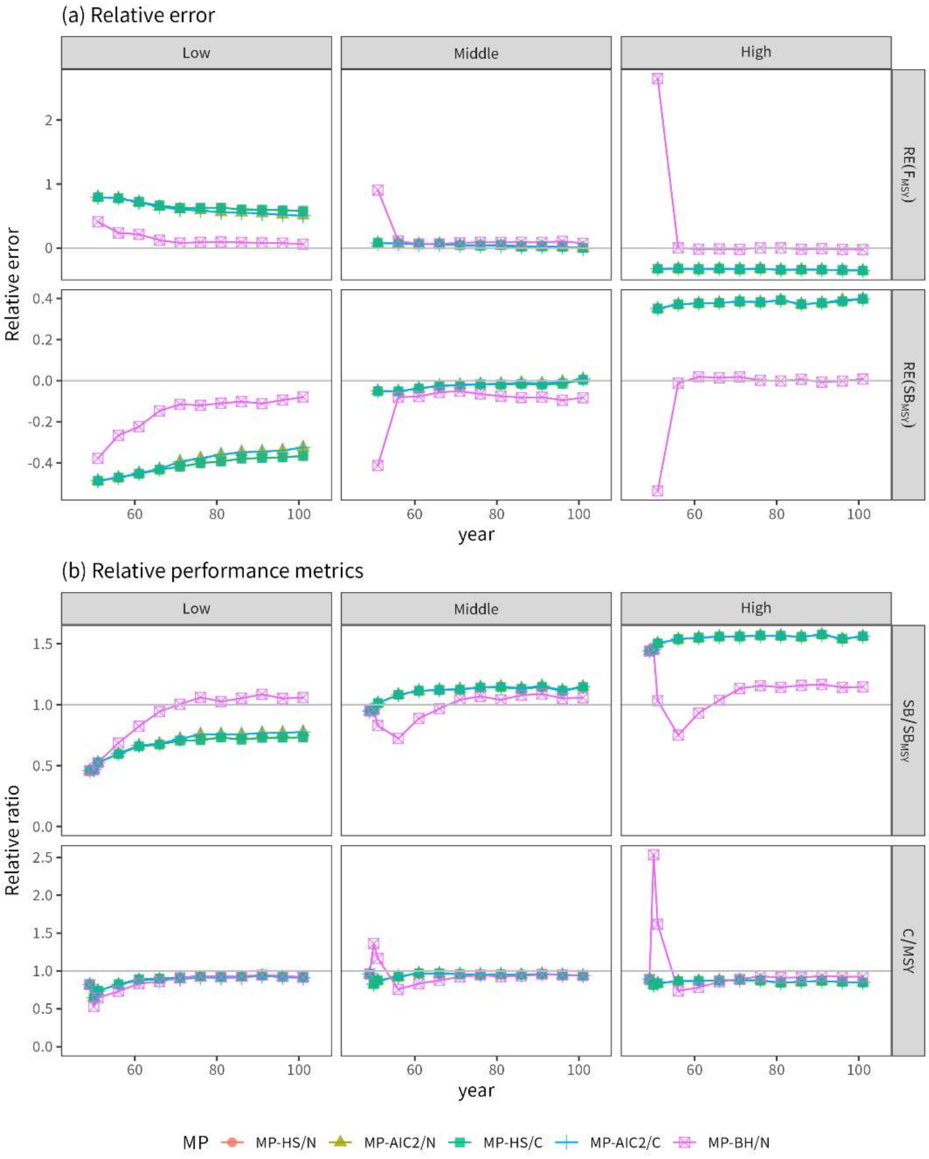


Fig. C5. Same as Fig. C4, but for the 3_MD species.

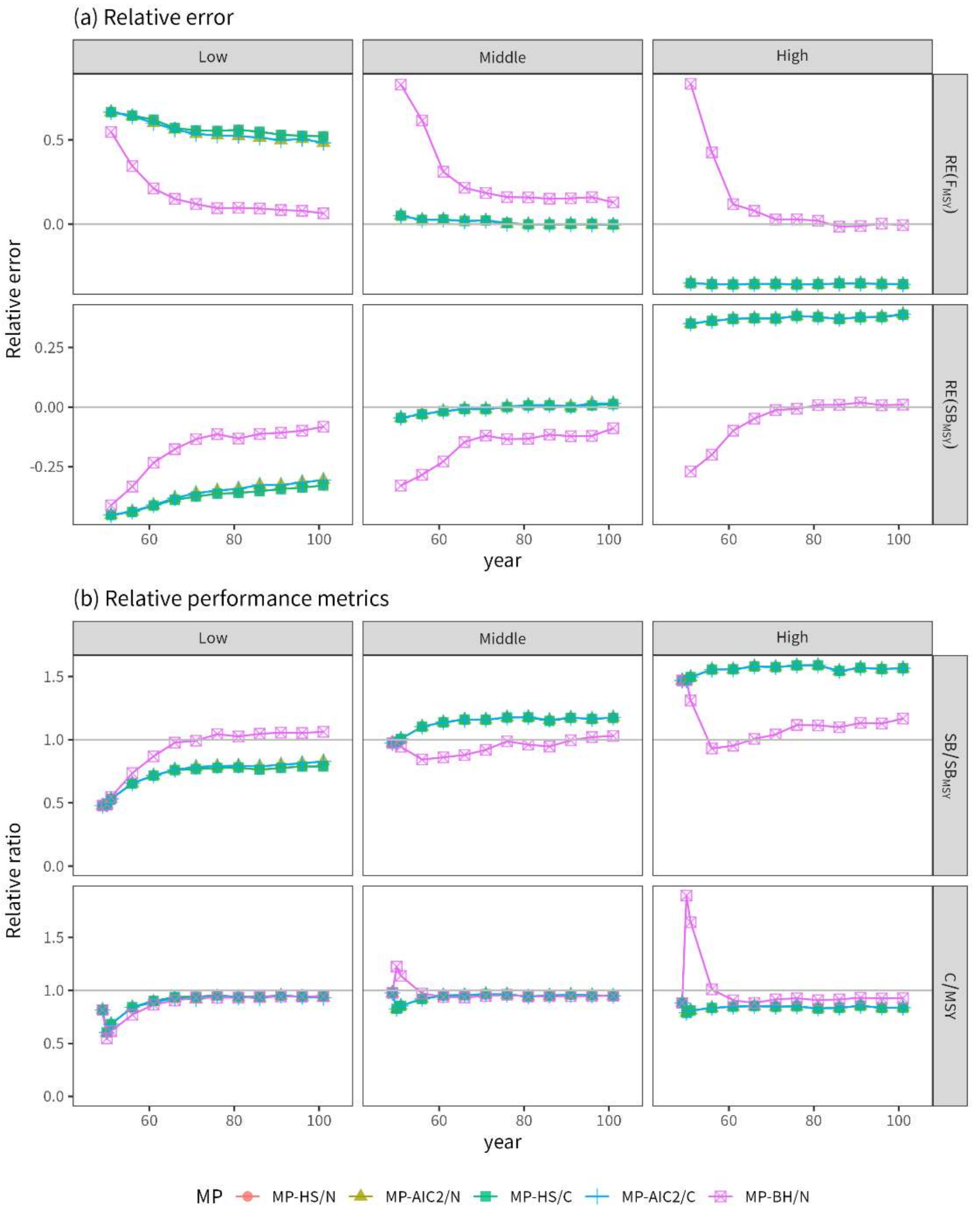


Fig. C6. Same as Fig. C4, but for the 4_LD species.

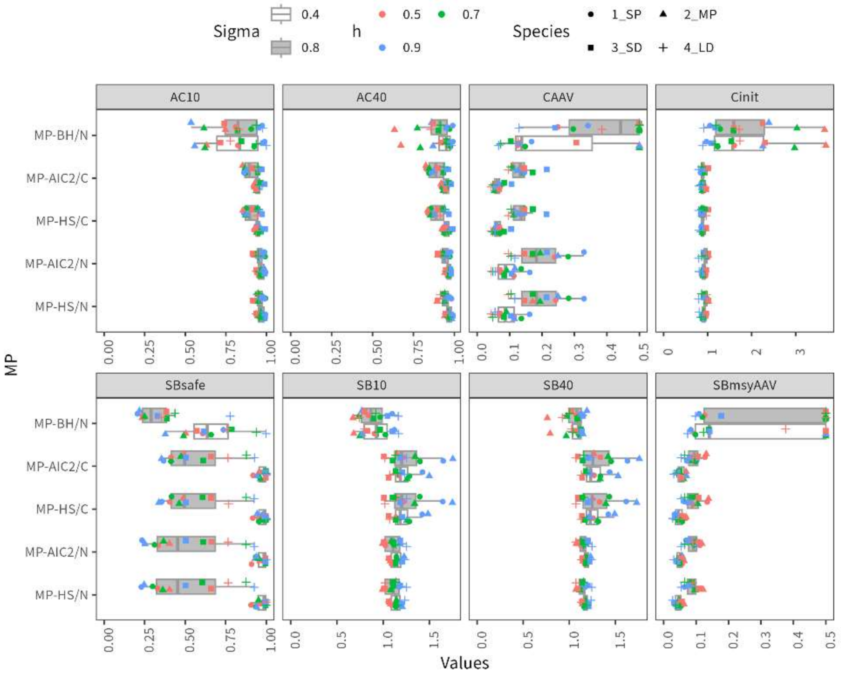


Fig. C7. Comparison among eight performance statistics yielded by five MPs under the “middle” scenario for all examined parameters and species. The maximum value of the y-axis for SBmsyAAV was set to 5; if it exceeded this value, it was plotted as 5.

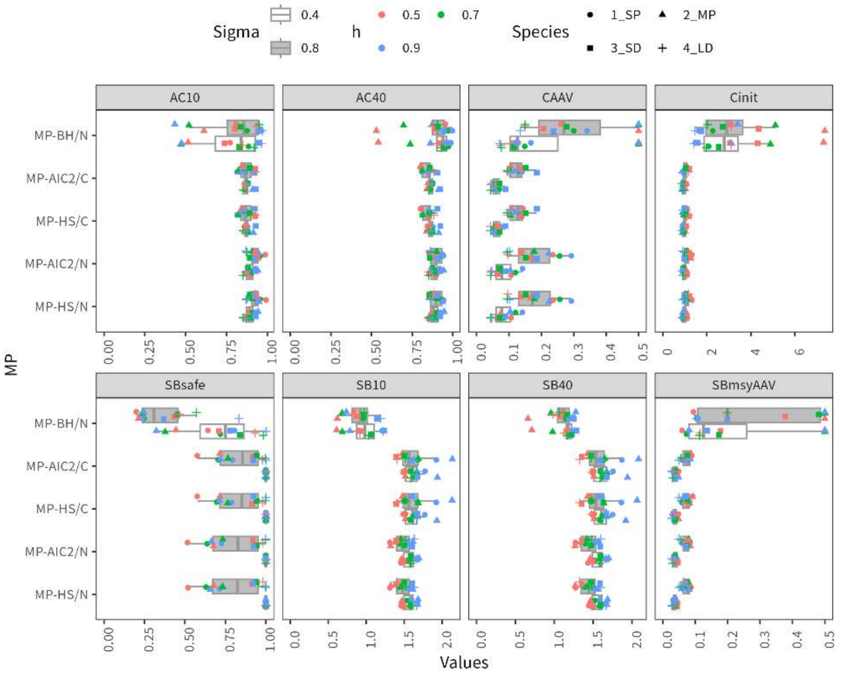


Fig. C8. Same as Fig. C7, but for the “high” scenario.

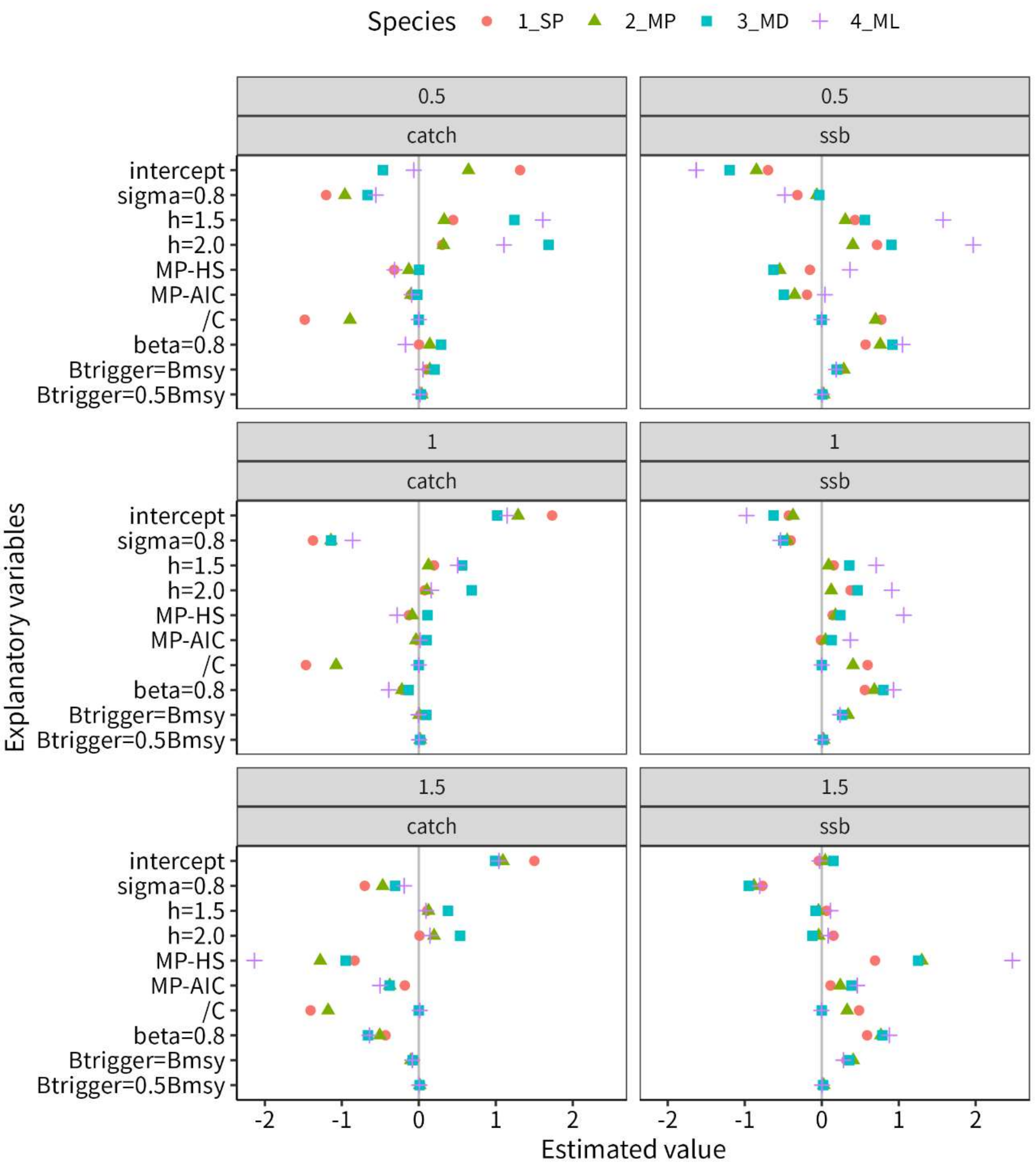


Fig. C9. Coefficients estimated for the GLM analysis under the sensitivity analysis when RI was used as the true SRR.

Fig. D. RE($F_{MSY}$), RE($SB_{MSY}$), $SB/SB_{MSY}$, and C/MSY under1_SP, Low, σ = 0.4

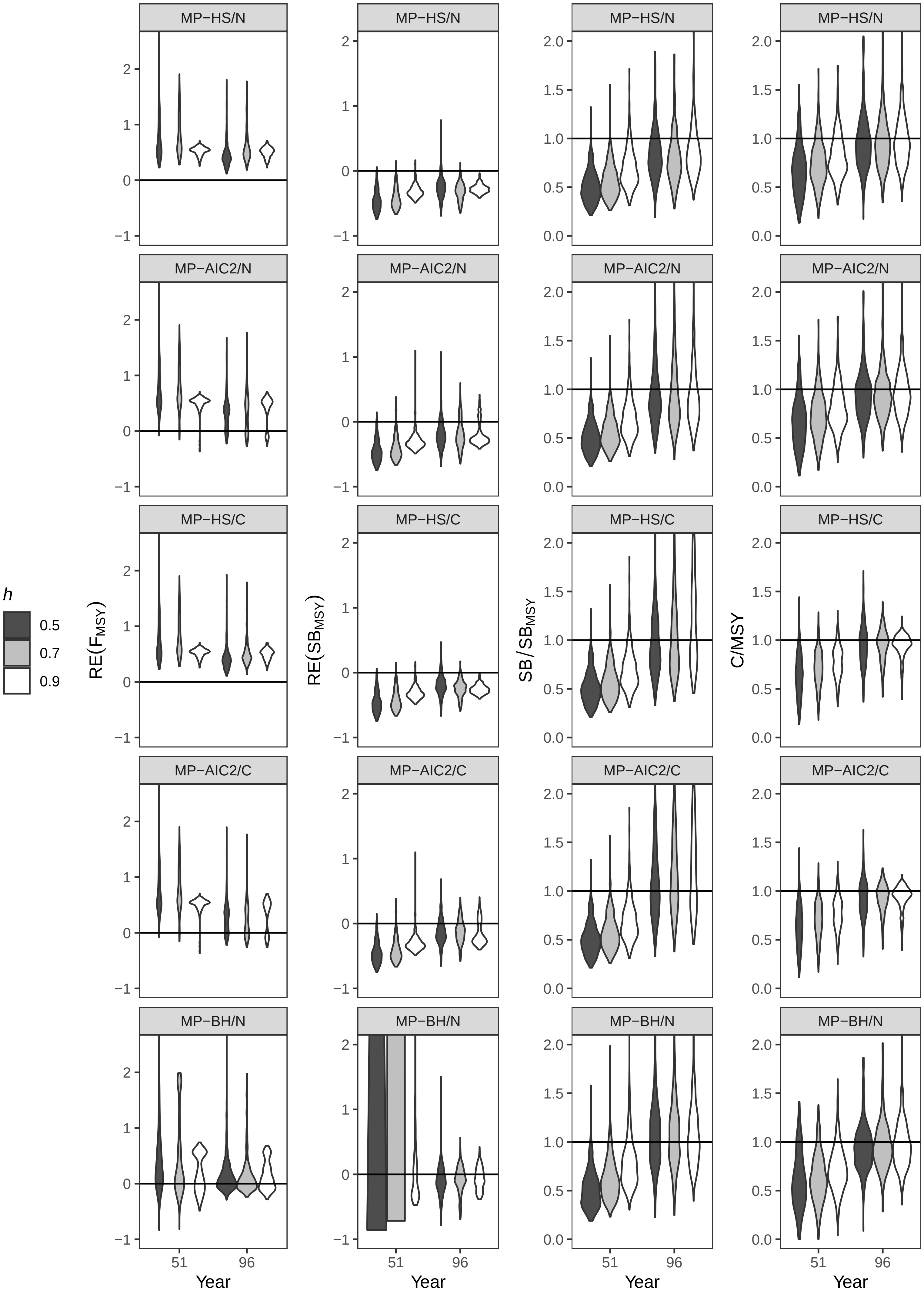

Fig. D. RE($F_{MSY}$), RE($SB_{MSY}$), SB/$SB_{MSY}$, and C/MSY under1_SP, Low, σ = 0.8

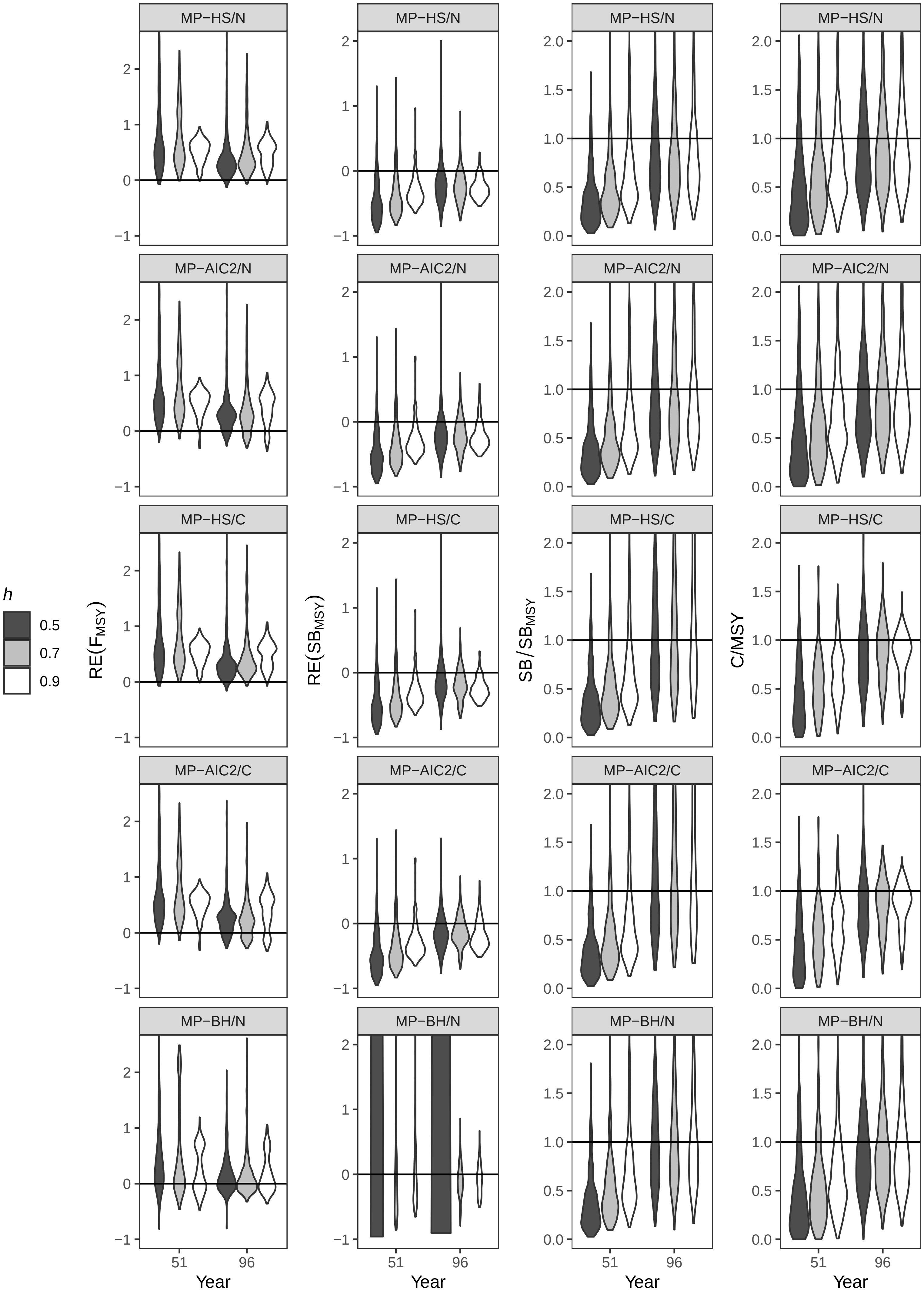

Fig. D. RE($F_{MSY}$), RE($SB_{MSY}$), SB/$SB_{MSY}$, and C/MSY under1_SP, Middle, σ = 0.4

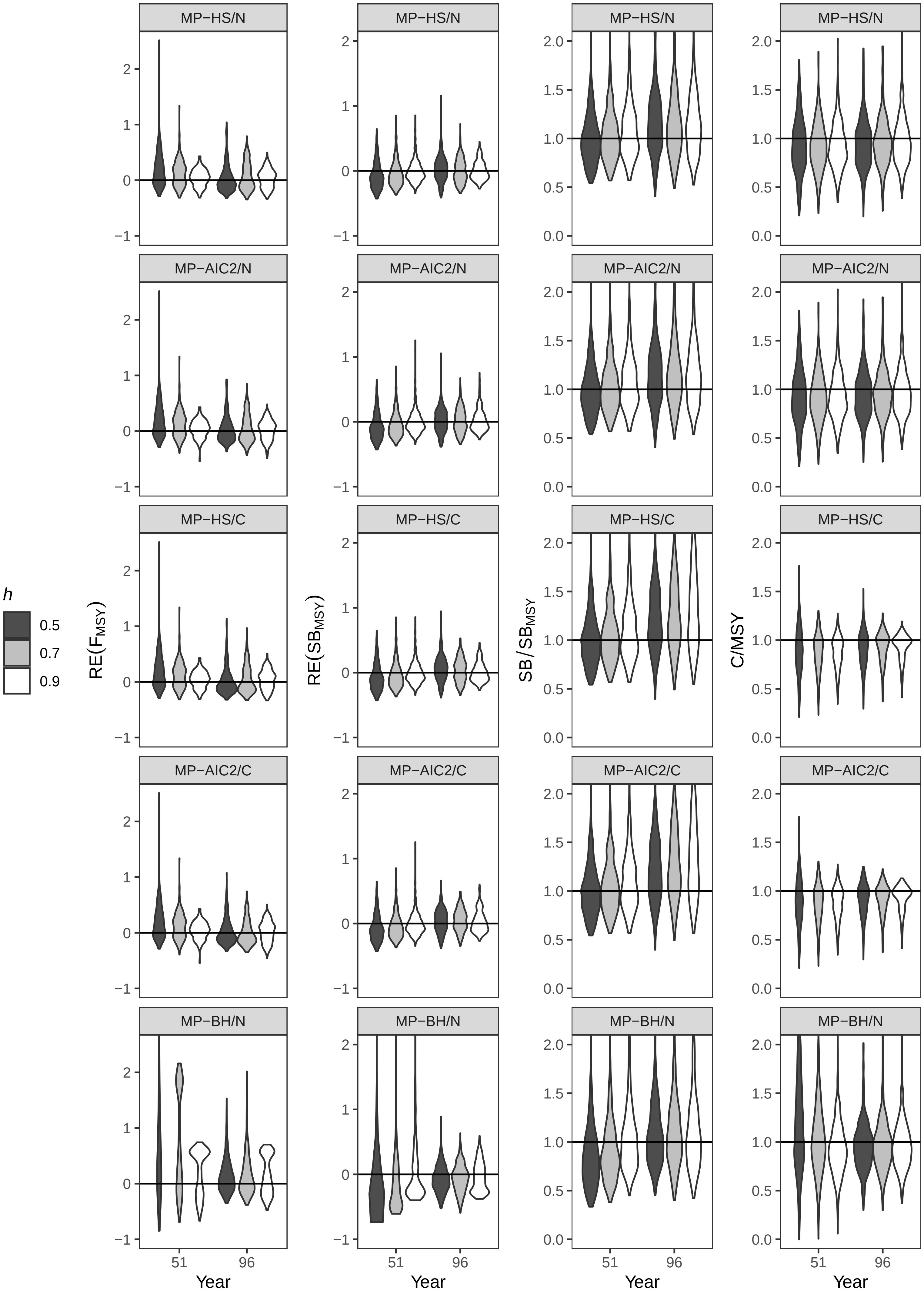

Fig. D. RE($F_{MSY}$), RE($SB_{MSY}$), $SB/SB_{MSY}$, and C/MSY under1_SP, Middle, σ = 0.8

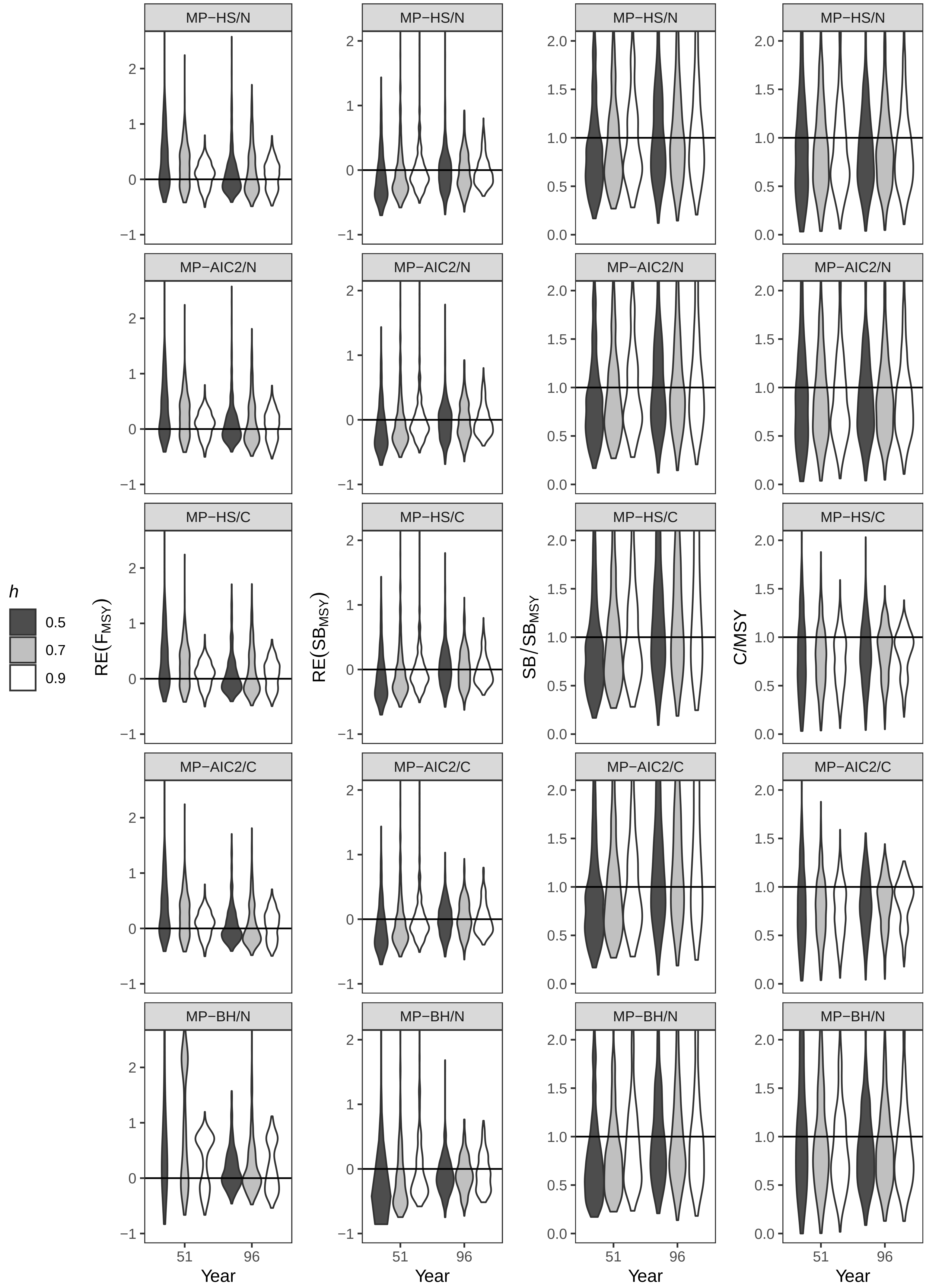

Fig. D. RE($F_{MSY}$), RE($SB_{MSY}$), $SB/SB_{MSY}$, and C/MSY under1_SP, High, σ = 0.4

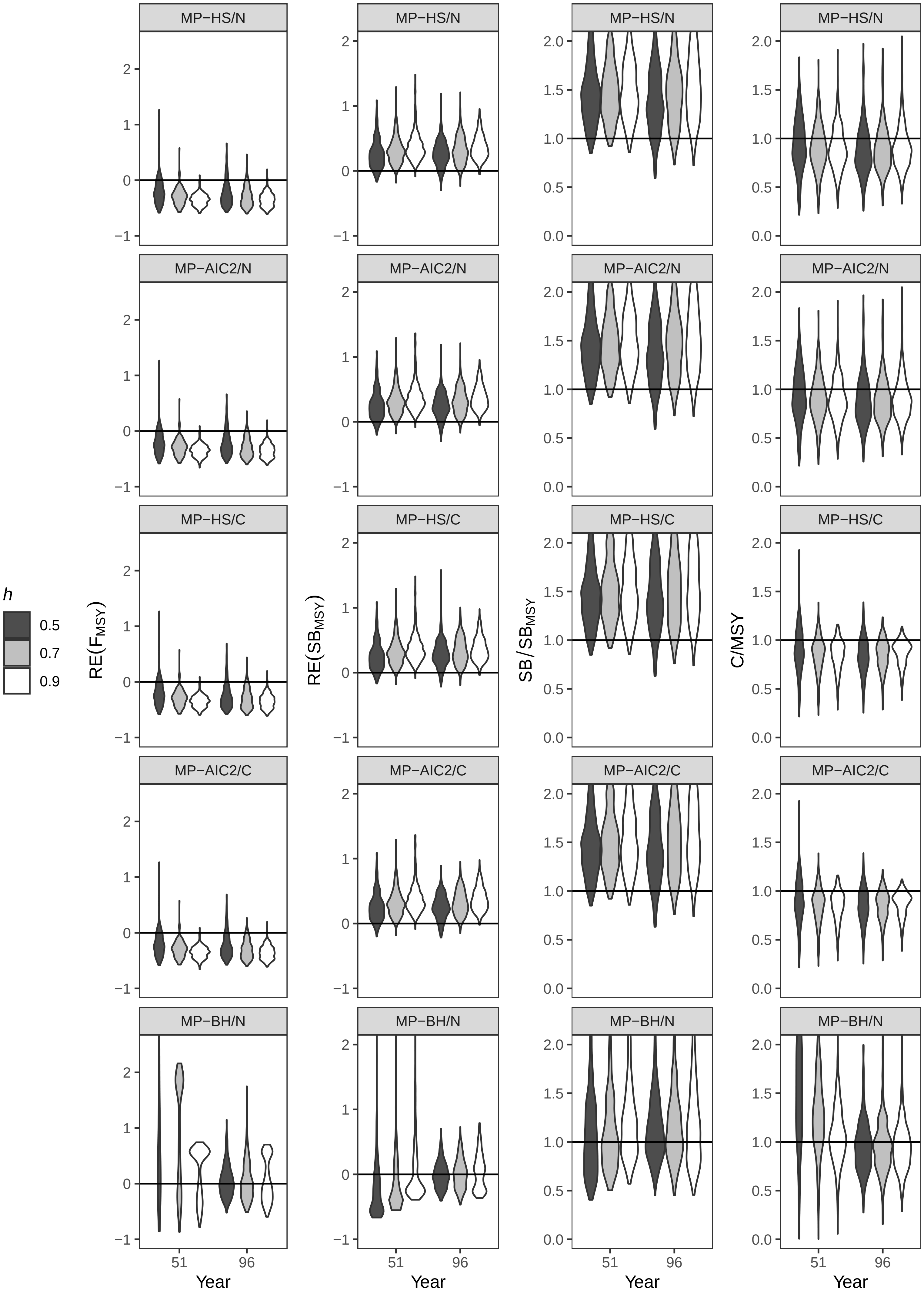

Fig. D. RE($F_{MSY}$), RE($SB_{MSY}$), $SB/SB_{MSY}$, and C/MSY under1_SP, High, σ = 0.8

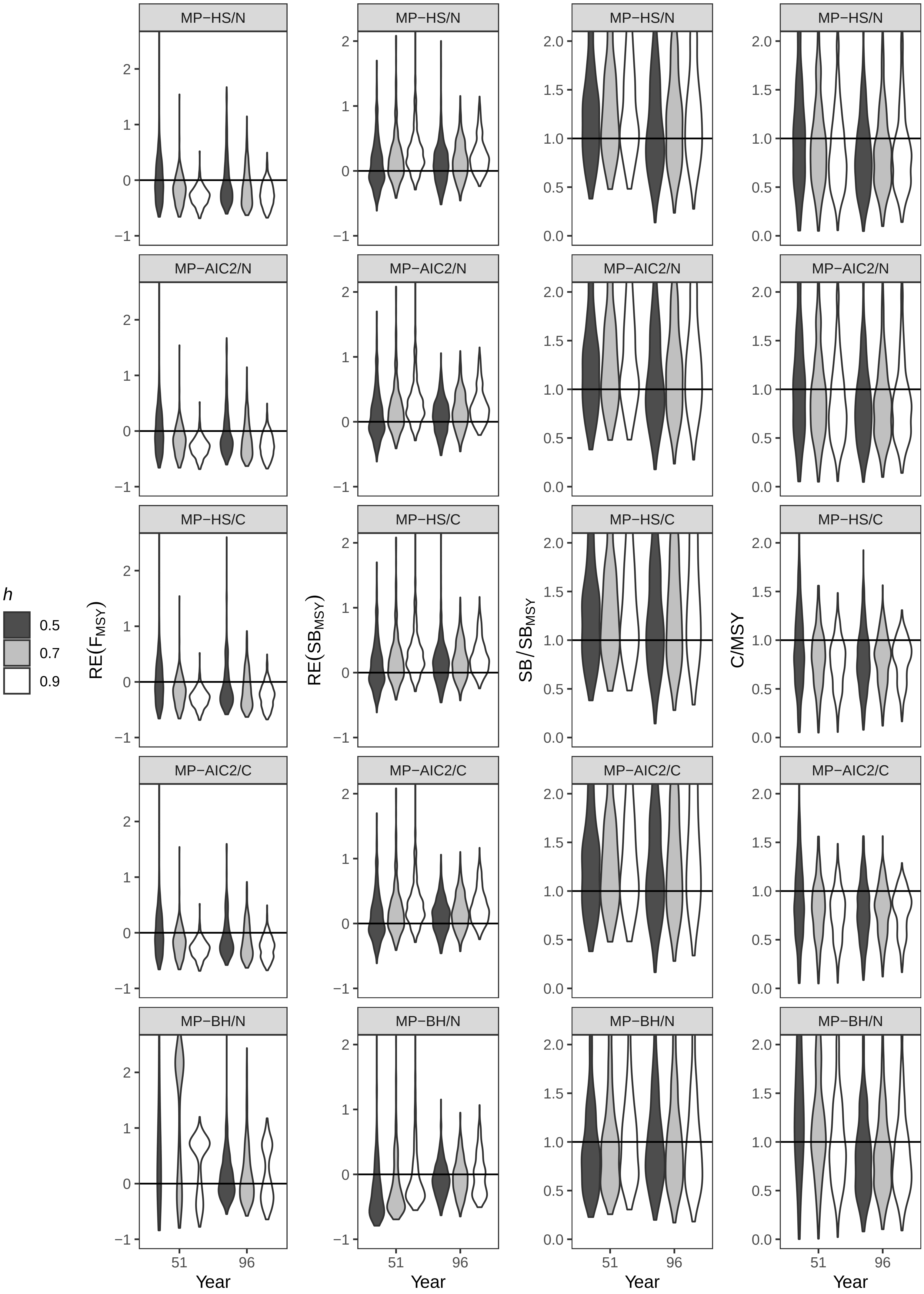

Fig. D. RE($F_{MSY}$), RE($SB_{MSY}$), $SB/SB_{MSY}$, and C/MSY under2_MP, Low, σ = 0.4

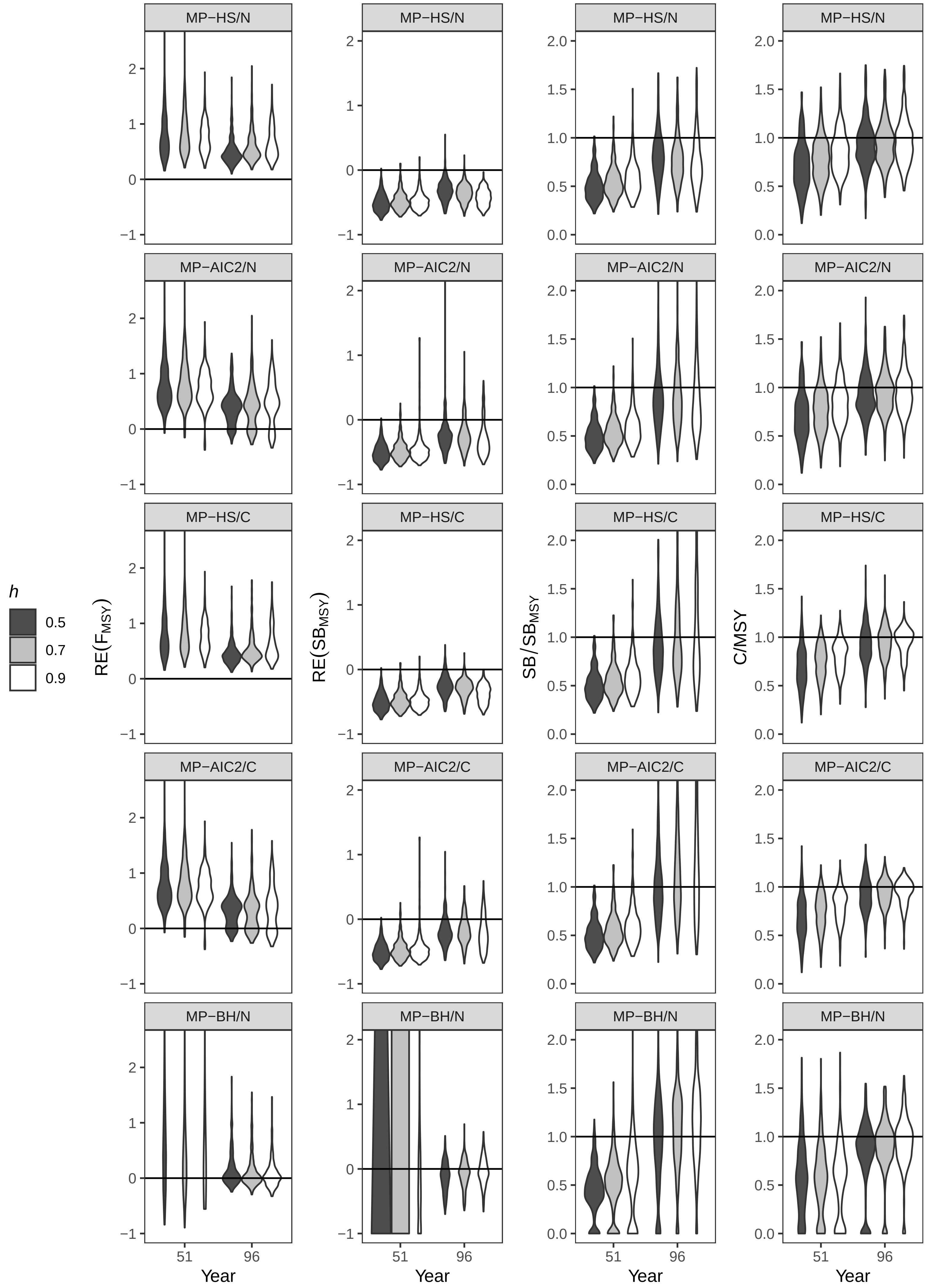

Fig. D. RE($F_{MSY}$), RE($SB_{MSY}$), $SB/SB_{MSY}$, and C/MSY under2_MP, Low, σ = 0.8

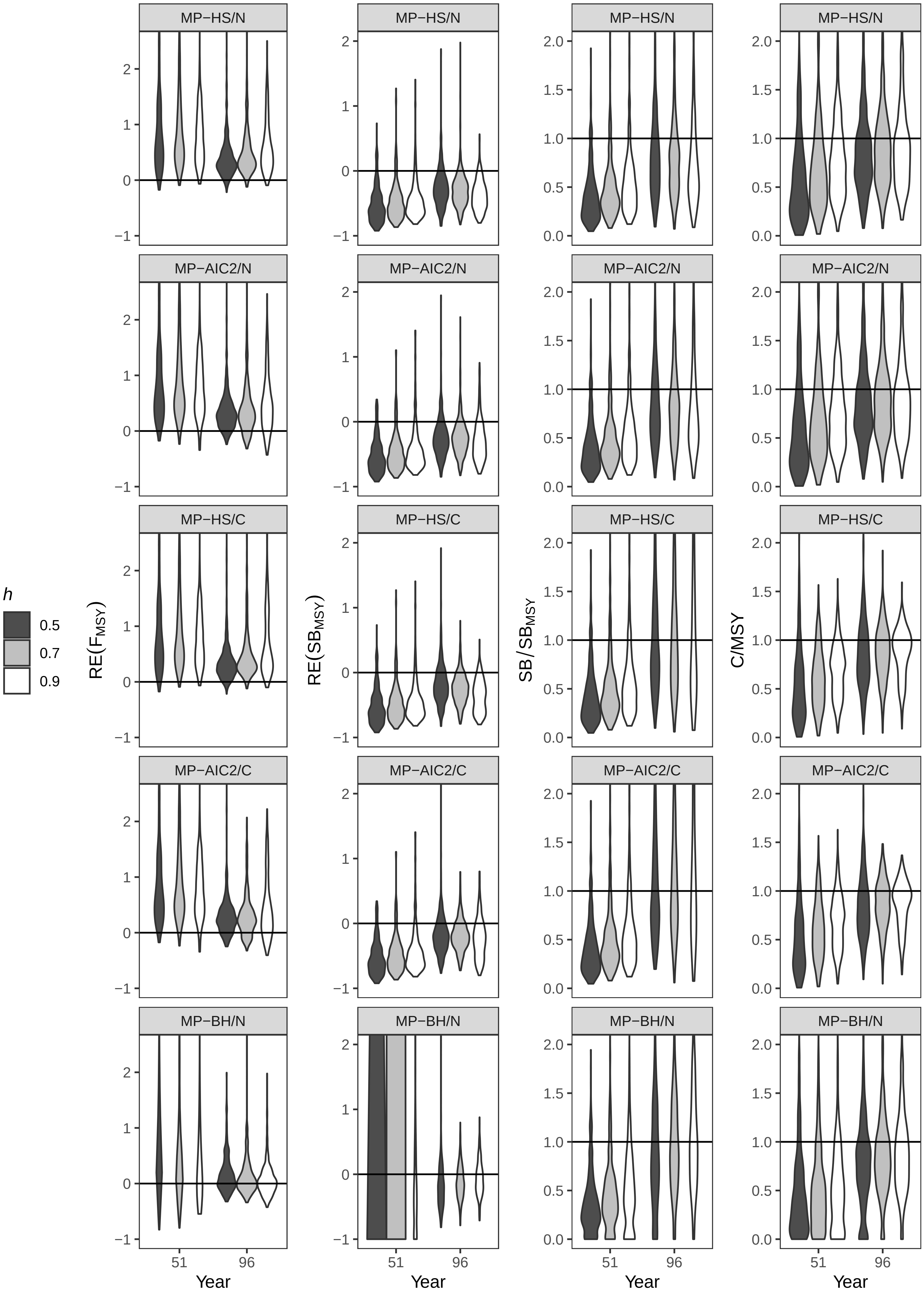

Fig. D. RE($F_{MSY}$), RE($SB_{MSY}$), SB/$SB_{MSY}$, and C/MSY under2_MP, Middle, σ = 0.4

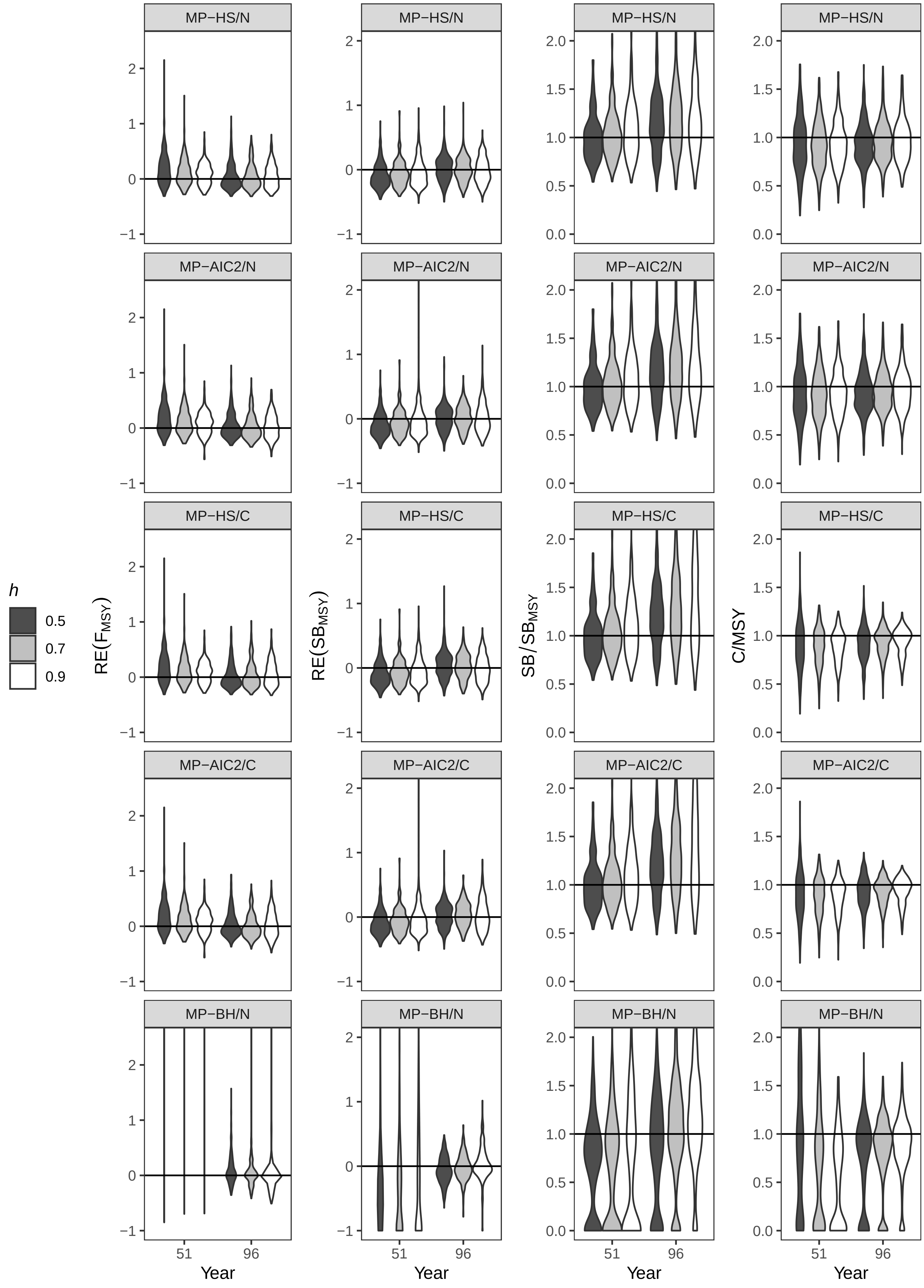

Fig. D. $RE(F_{MSY})$, $RE(SB_{MSY})$, $SB/SB_{MSY}$, and C/MSY under2_MP, Middle, $\sigma$ = 0.8

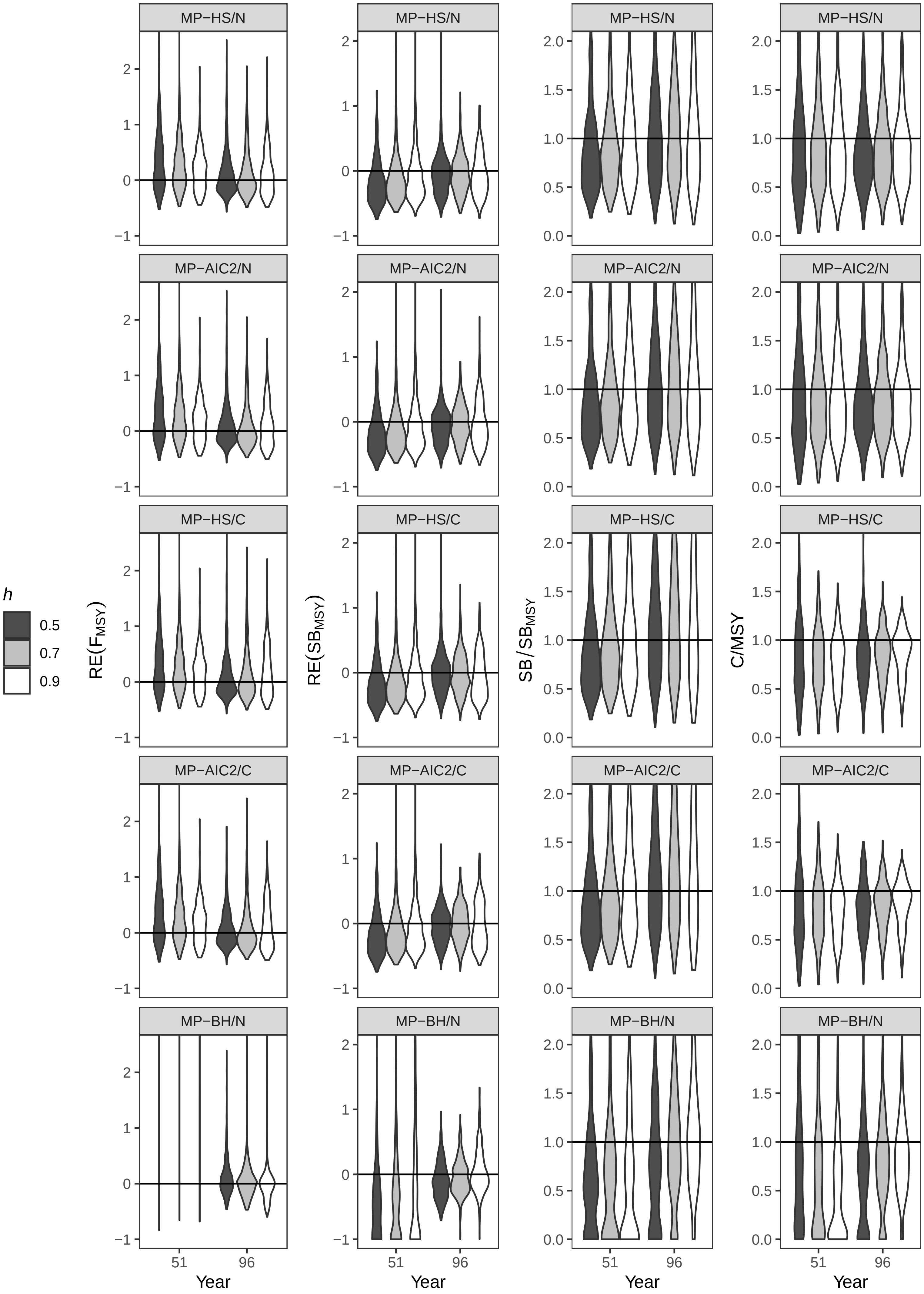

Fig. D. RE($F_{MSY}$), RE($SB_{MSY}$), $SB/SB_{MSY}$, and C/MSY under2_MP, High, $\sigma$ = 0.4

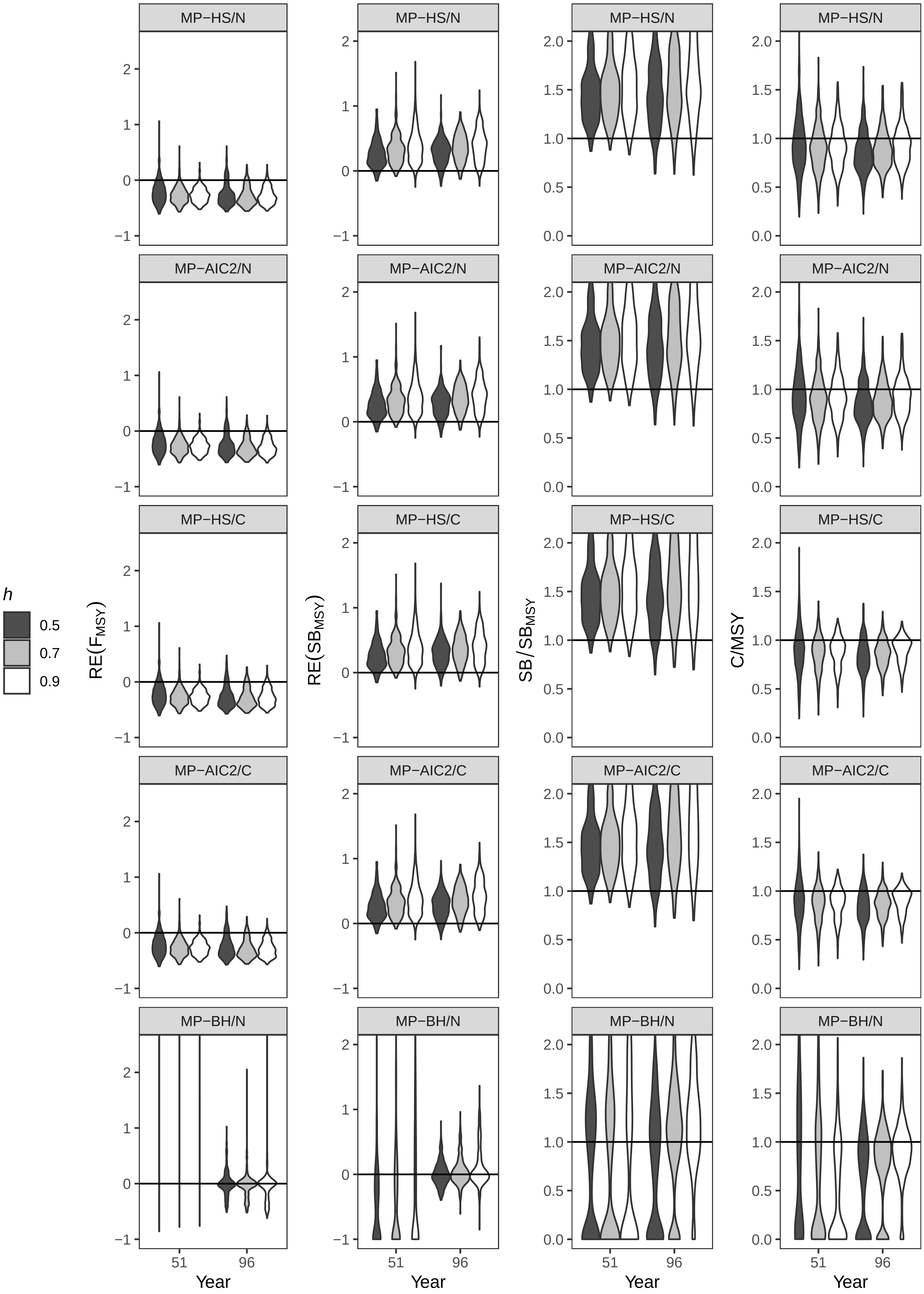

Fig. D. RE($F_{MSY}$), RE($SB_{MSY}$), $SB/SB_{MSY}$, and C/MSY under2_MP, High, $\sigma$ = 0.8

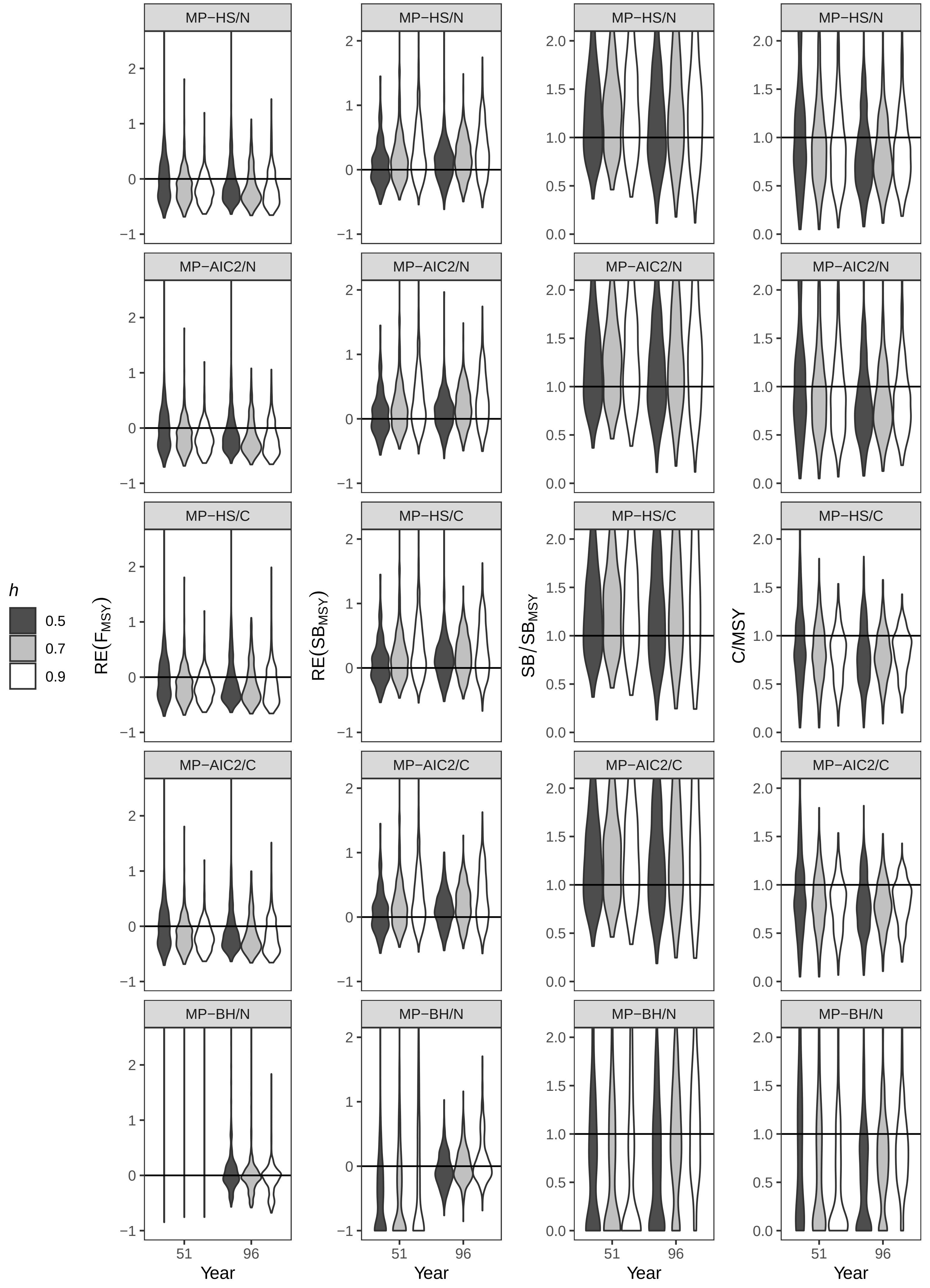

Fig. D. RE($F_{MSY}$), RE($SB_{MSY}$), $SB/SB_{MSY}$, and C/MSY under3_MD, Low, σ = 0.4

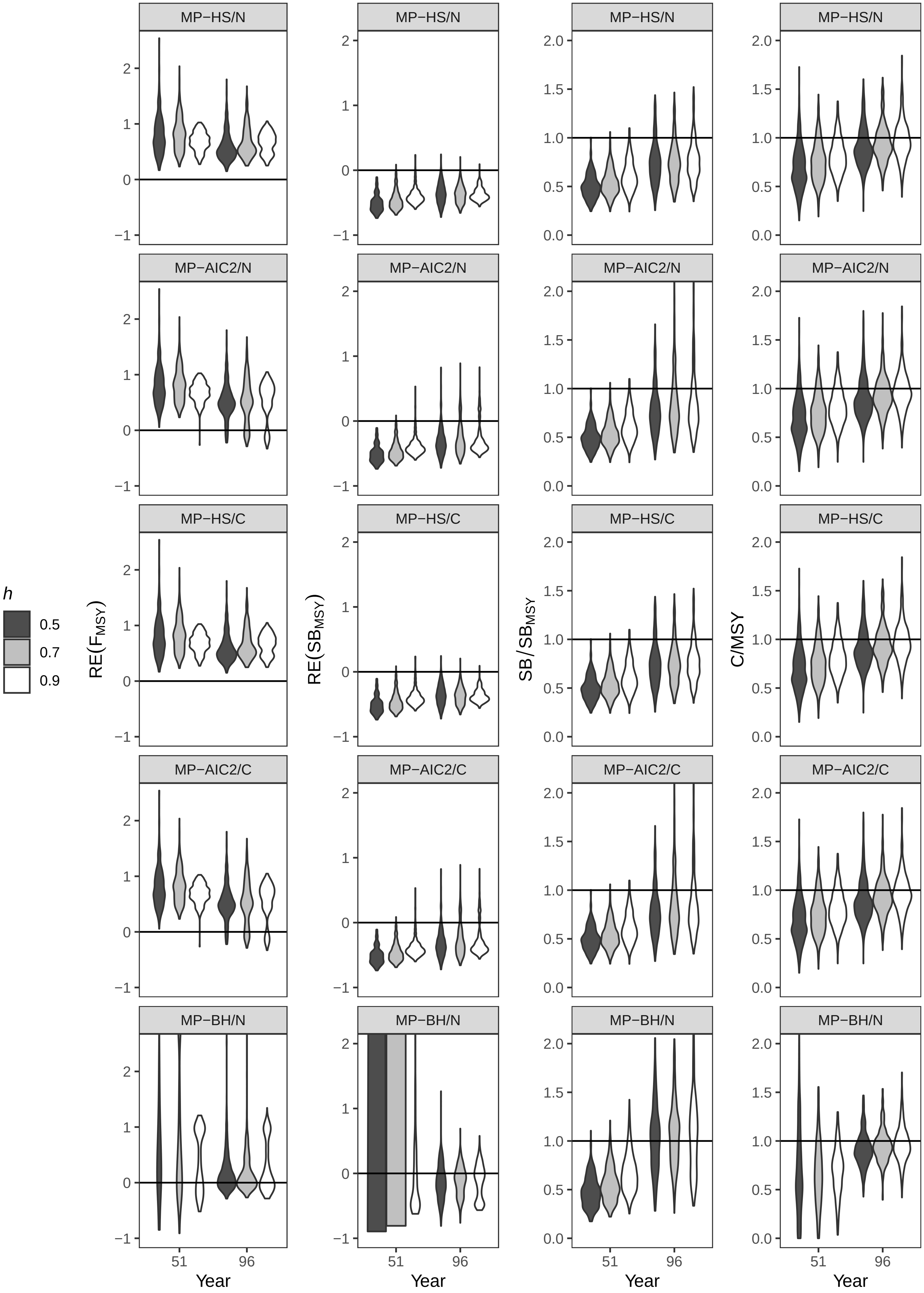

Fig. D. RE($F_{MSY}$), RE($SB_{MSY}$), SB/$SB_{MSY}$, and C/MSY under3_MD, Low, σ = 0.8

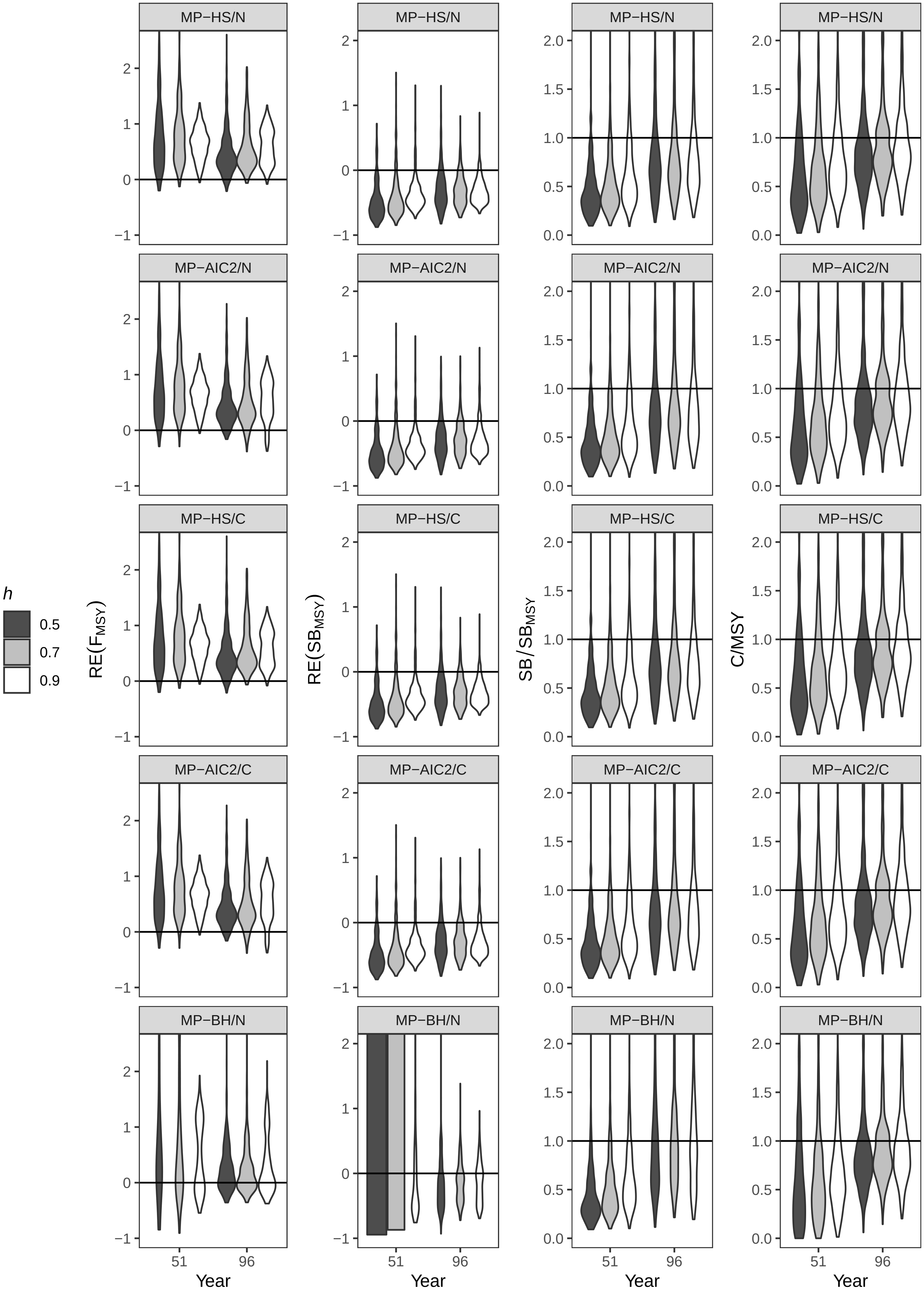

Fig. D. RE($F_{MSY}$), RE($SB_{MSY}$), $SB/SB_{MSY}$, and C/MSY under3_MD, Middle, $\sigma$ = 0.4

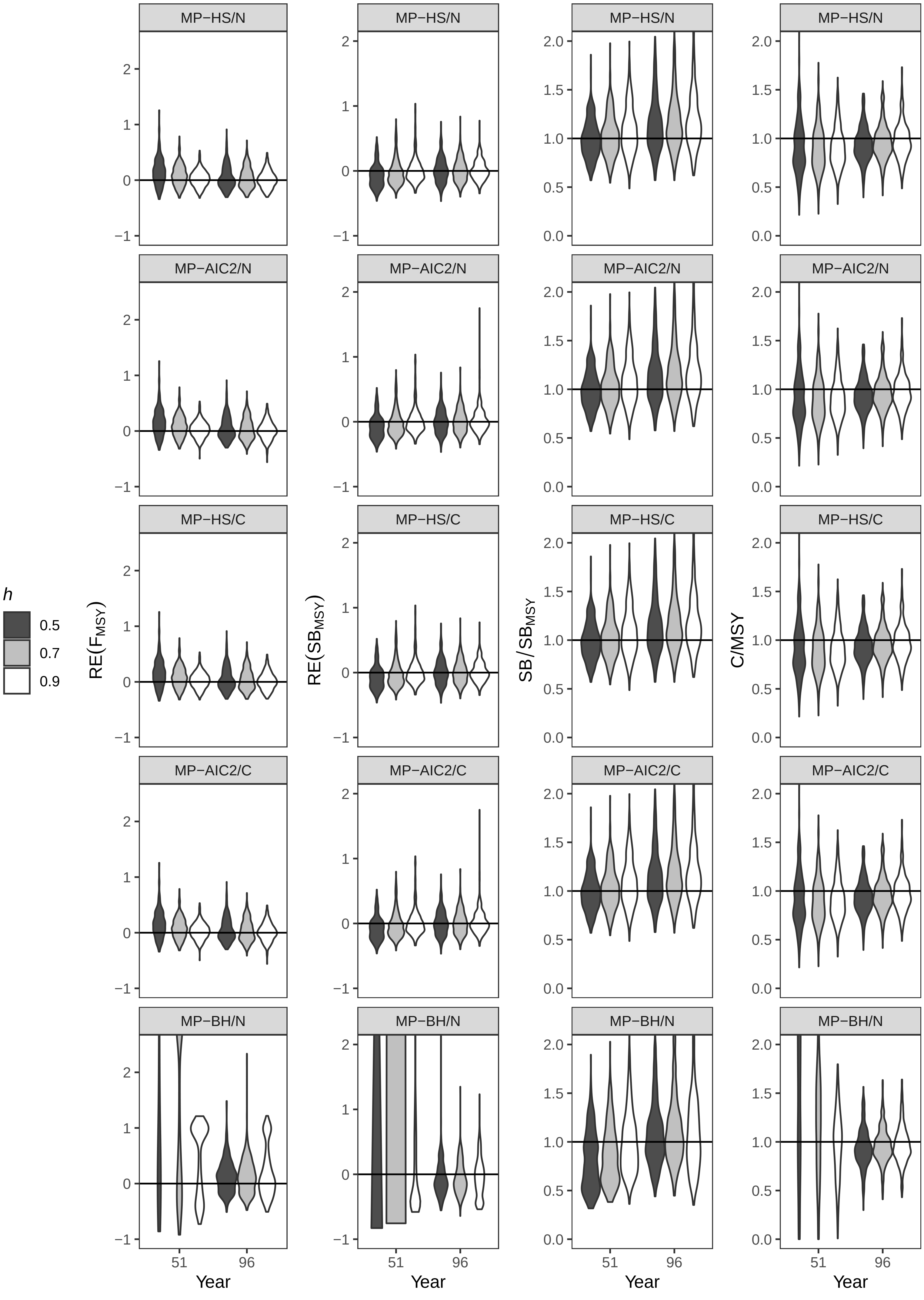

Fig. D. RE($F_{MSY}$), RE($SB_{MSY}$), $SB/SB_{MSY}$, and C/MSY under3_MD, Middle, $\sigma$ = 0.8

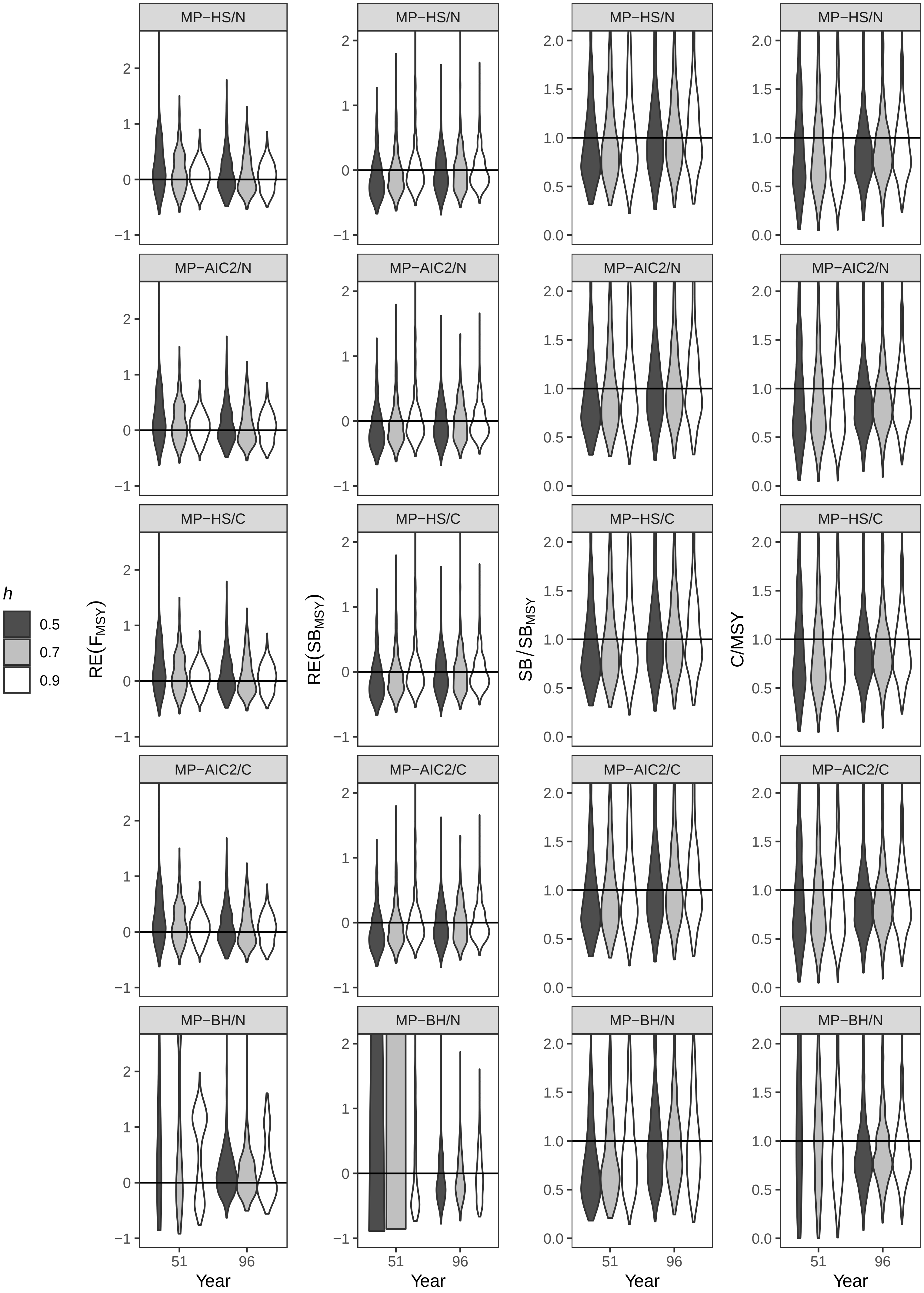

Fig. D. RE($F_{MSY}$), RE($SB_{MSY}$), SB/$SB_{MSY}$, and C/MSY under3_MD, High, σ = 0.4

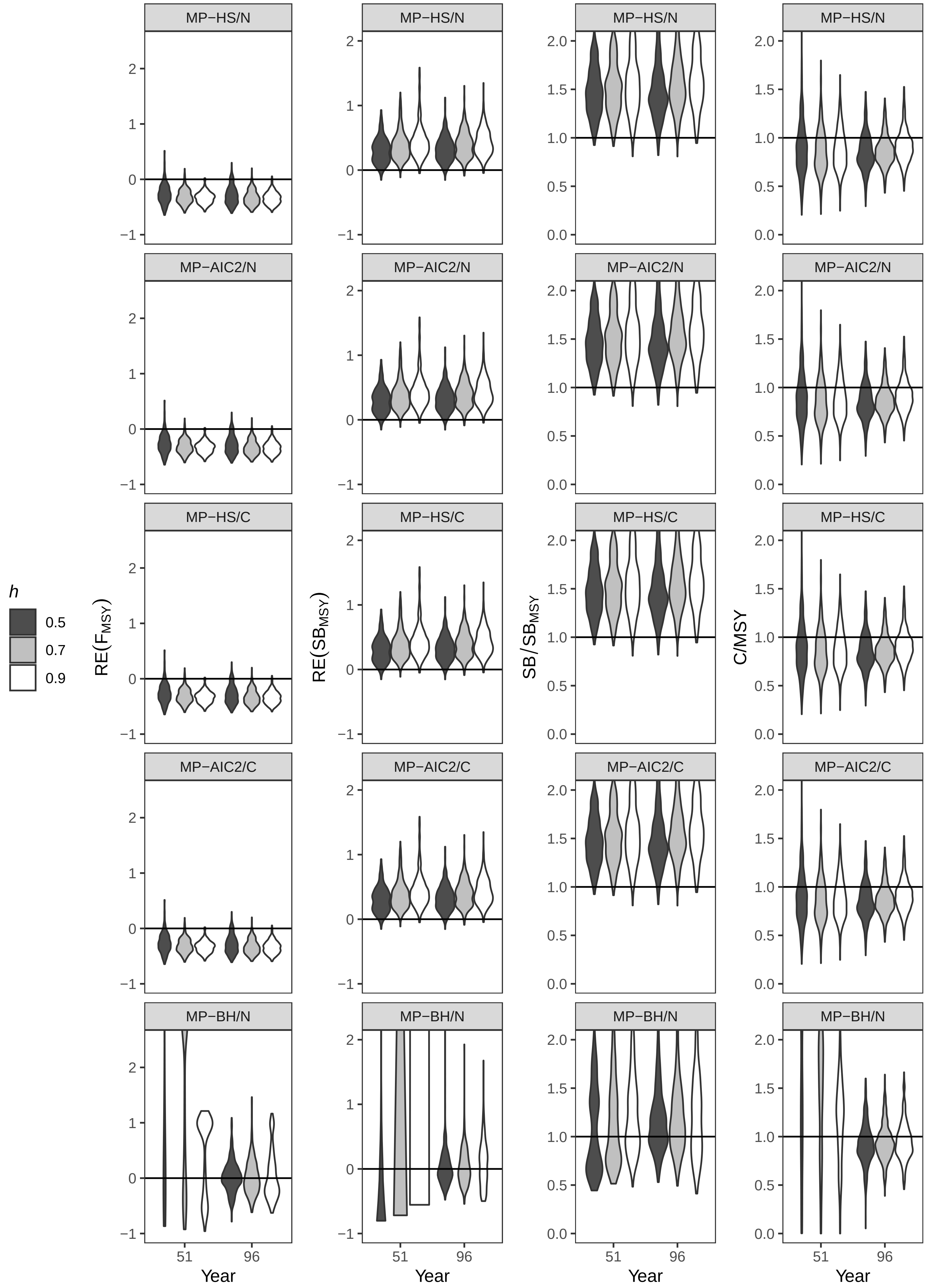

Fig. D. RE($F_{MSY}$), RE($SB_{MSY}$), $SB/SB_{MSY}$, and C/MSY under3_MD, High, $\sigma$ = 0.8

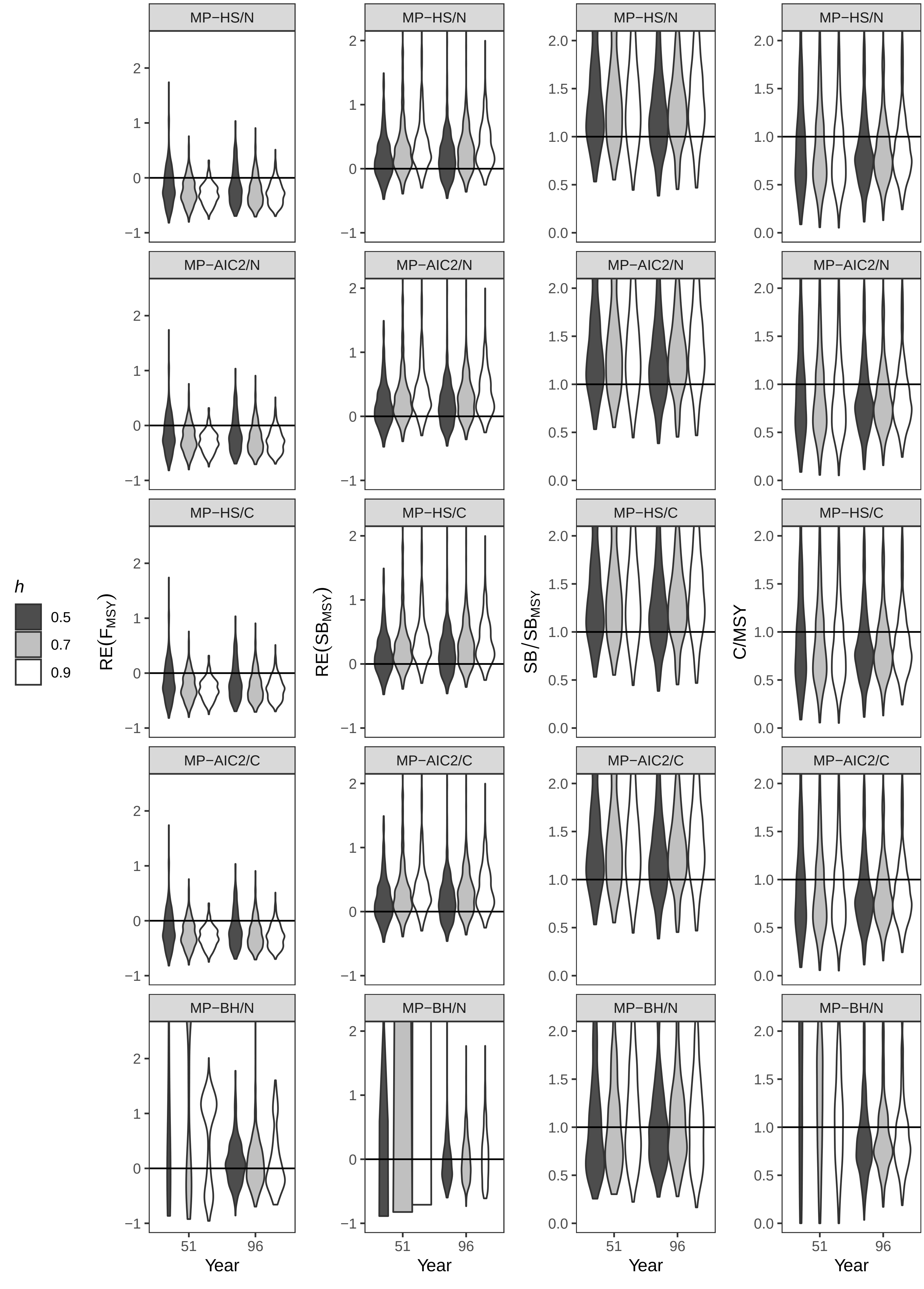

Fig. D. RE($F_{MSY}$), RE($SB_{MSY}$), $SB/SB_{MSY}$, and C/MSY under4_LD, Low, $\sigma$ = 0.4

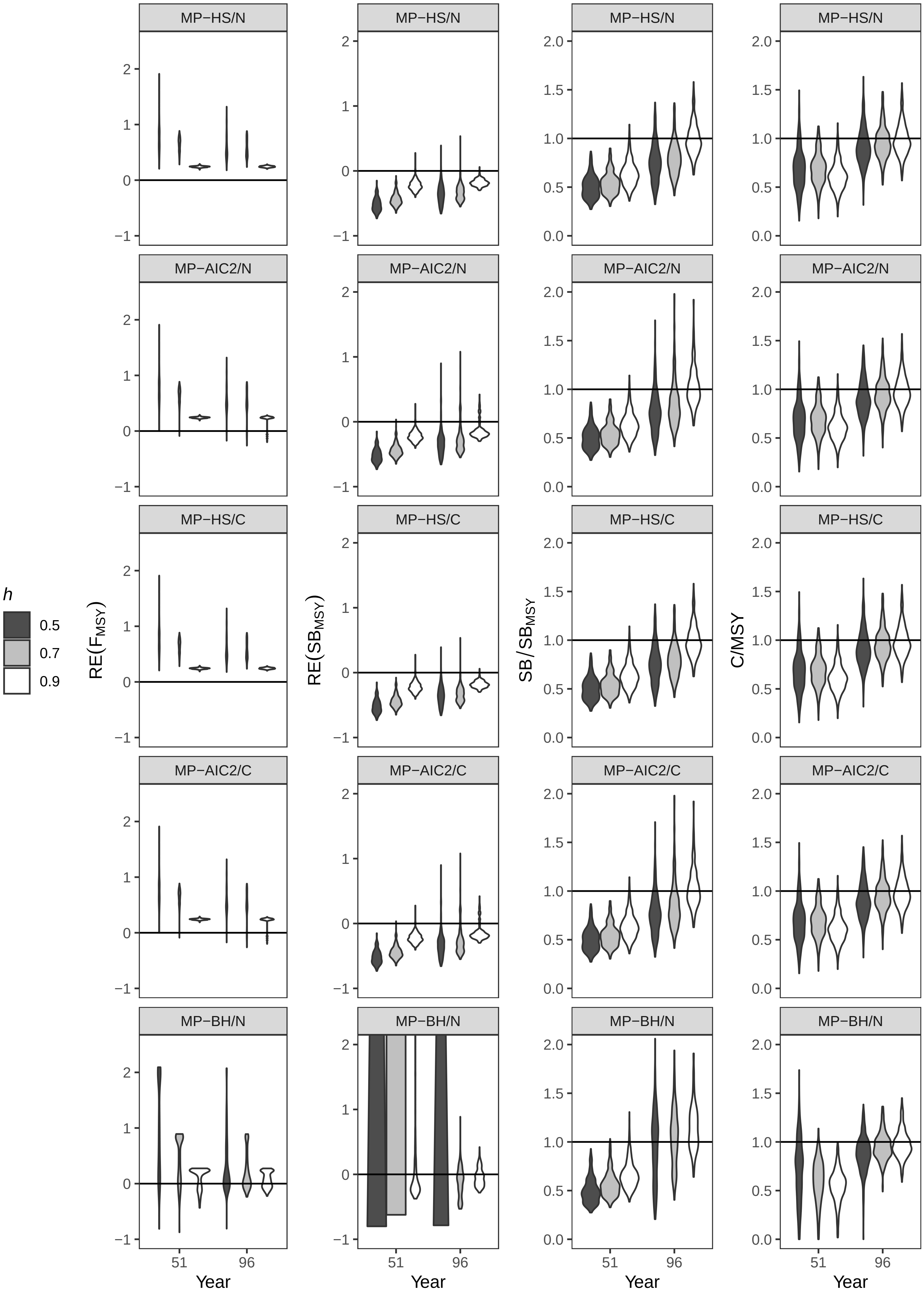

Fig. D. RE($F_{MSY}$), RE($SB_{MSY}$), $SB/SB_{MSY}$, and C/MSY under4_LD, Low, σ = 0.8

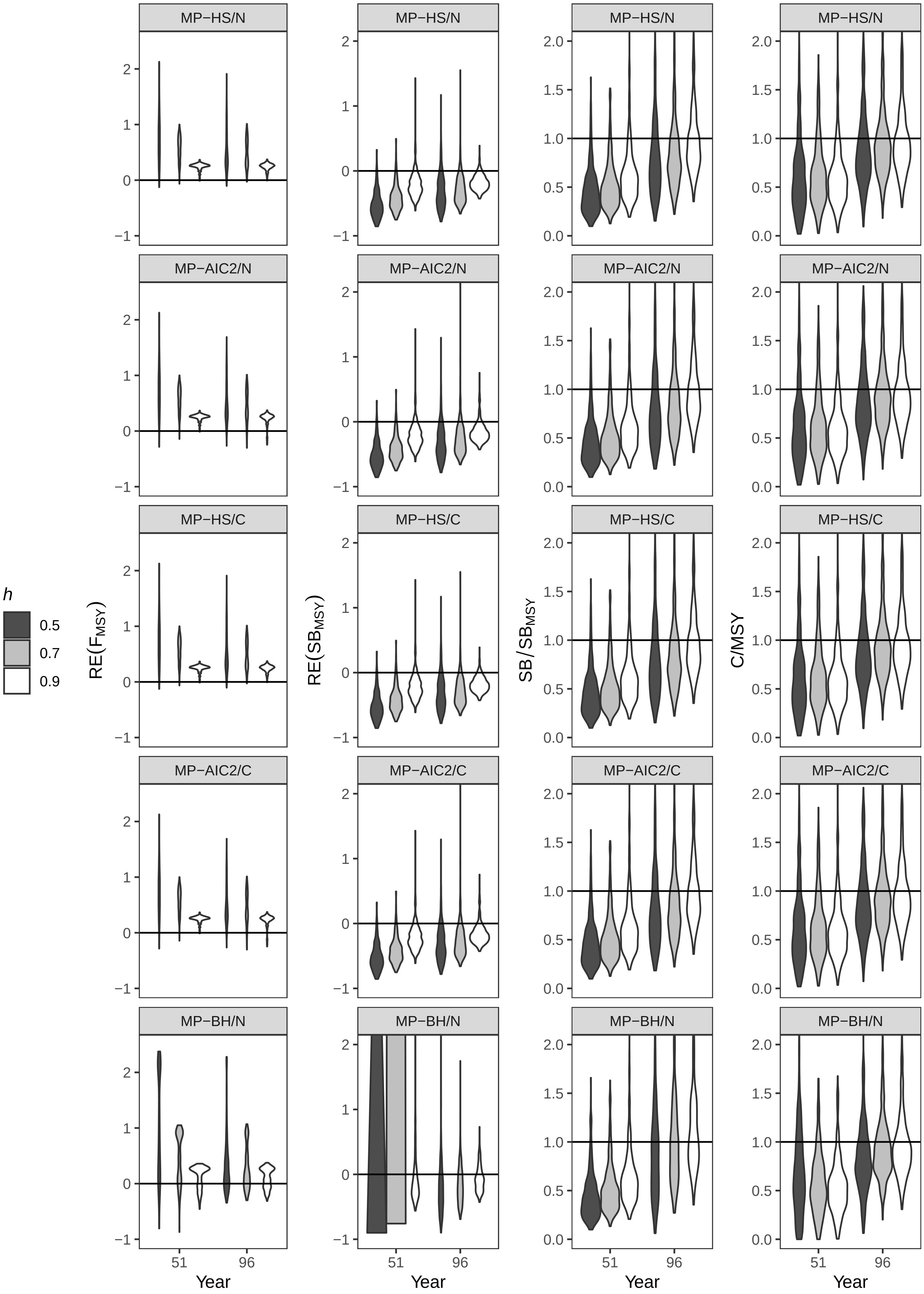

Fig. D. RE($F_{MSY}$), RE($SB_{MSY}$), SB/$SB_{MSY}$, and C/MSY under4_LD, Middle, σ = 0.4

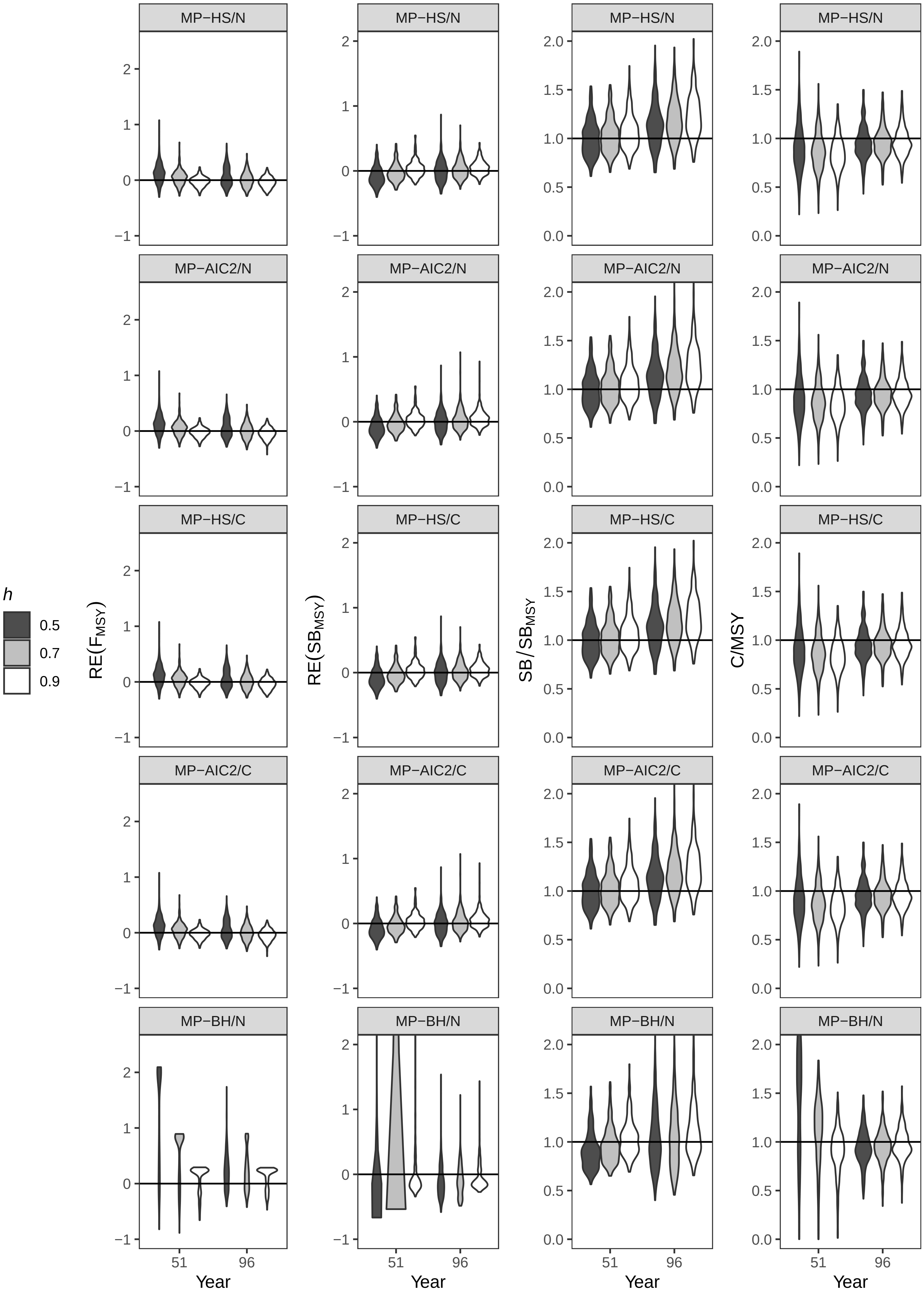

Fig. D. $RE(F_{MSY})$, $RE(SB_{MSY})$, $SB/SB_{MSY}$, and C/MSY under4_LD, Middle, $\sigma = 0.8$

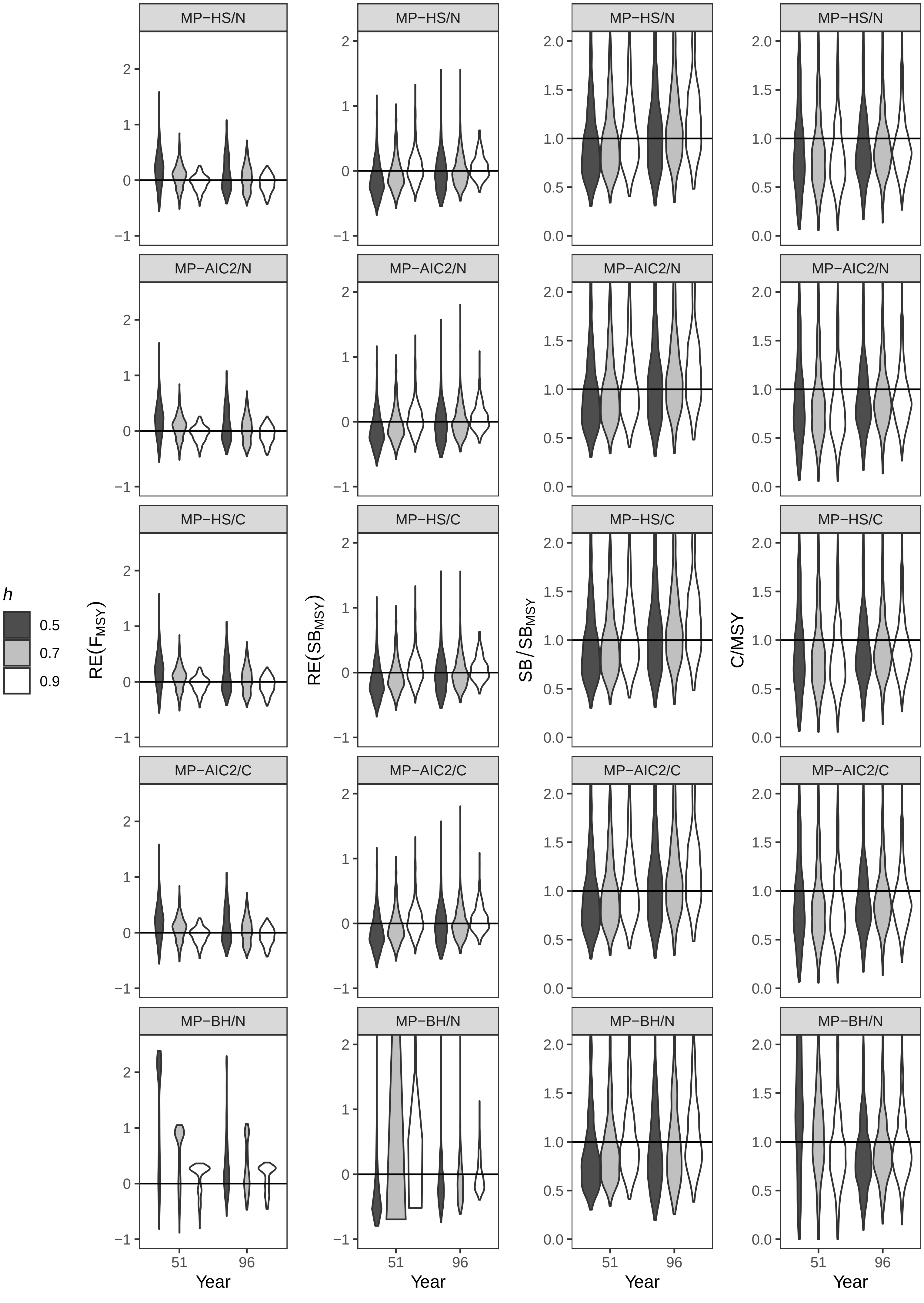

Fig. D. $RE(F_{MSY})$, $RE(SB_{MSY})$, $SB/SB_{MSY}$, and C/MSY under4_LD, High, $\sigma = 0.4$

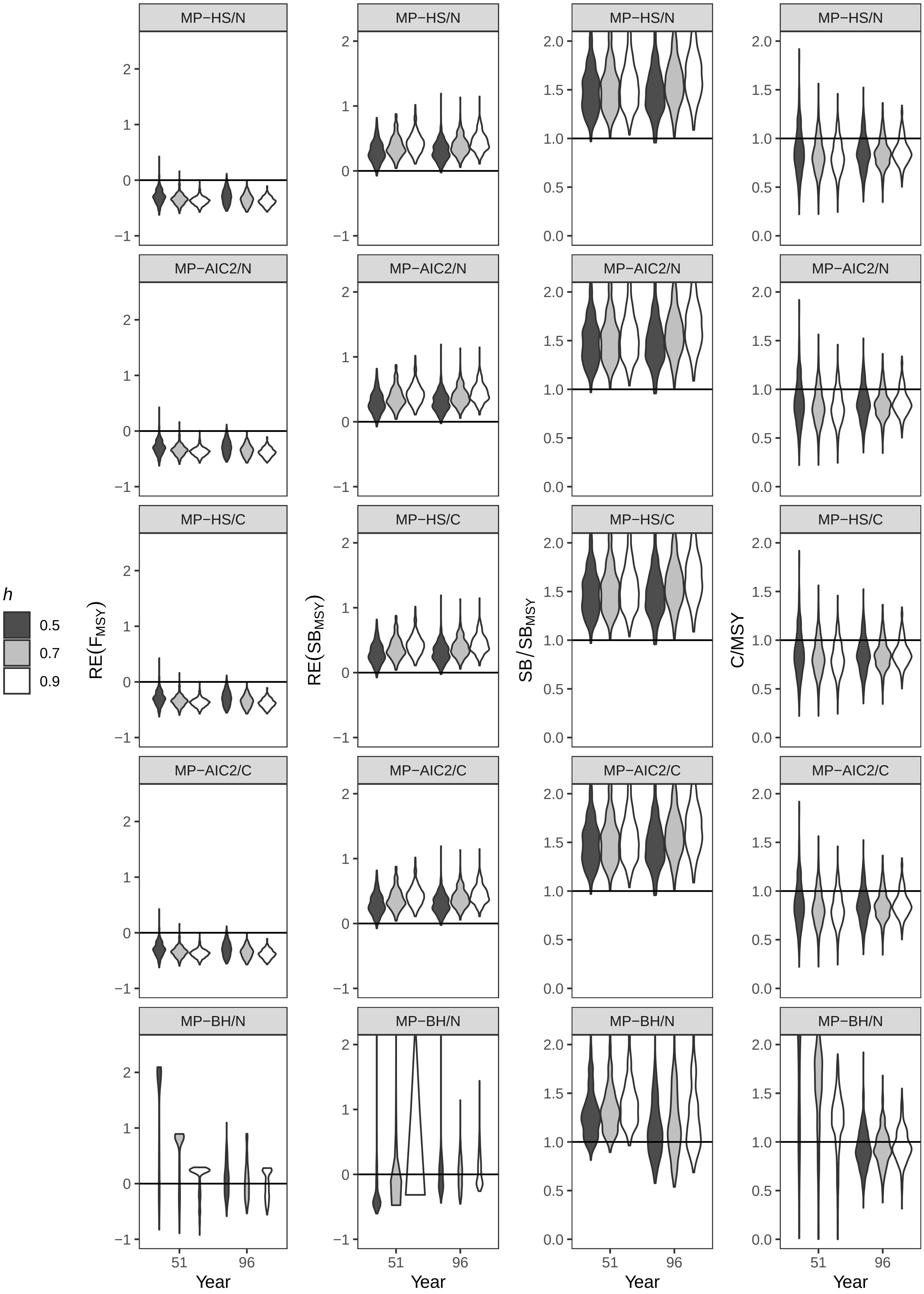

Fig. D. RE($F_{MSY}$), RE($SB_{MSY}$), SB/$SB_{MSY}$, and C/MSY under4_LD, High, σ = 0.8

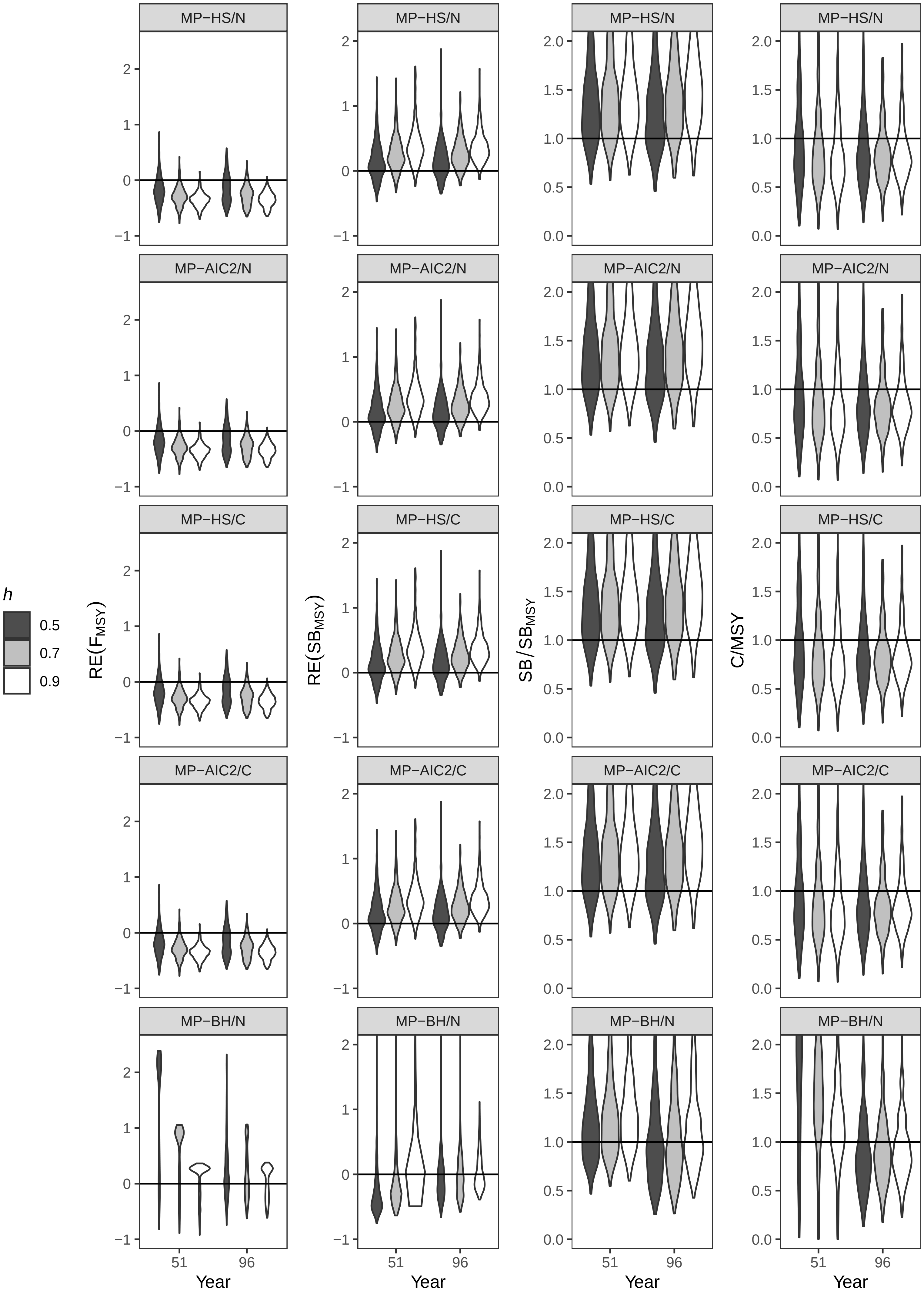
